# Histone chaperone HIRA regulates adiponectin expression and obesity-associated adipose expansion by facilitating Pol II pause release

**DOI:** 10.1101/2025.03.21.644577

**Authors:** Danyang Wan, Ji-Eun Lee, Young-Kwon Park, Guojia Xie, Susanna Maisto, Christabelle Agyapong, Keiko Ozato, Oksana Gavrilova, Kai Ge

## Abstract

Adipose tissue is essential for maintaining glucose and lipid homeostasis in mammals. However, epigenomic mechanisms underlying adipose tissue function remain largely unclear. Here, we identify the histone chaperon HIRA as a novel epigenomic regulator of adipose tissue function. Adipose tissue-specific knockout of *Hira* in mice impairs insulin sensitivity and restrains adipose tissue expansion during high-fat diet-induced obesity. Mechanistically, HIRA is required for the expression of *Adipoq*, encoding the adipokine adiponectin, and lipid metabolism genes in adipose tissue. Genomic mapping reveals that HIRA binds to promoters and enhancers of *Adipoq* and lipid metabolism genes in adipocytes. Acute HIRA depletion using the dTAG system, followed by nascent RNA-Seq and ChIP-Seq, demonstrates that while HIRA is largely dispensable for enhancer activation and coactivator binding, it promotes transcription of target genes by facilitating RNA polymerase II pause release and subsequent elongation likely independently of H3.3 deposition. Our findings uncover a novel mechanism by which HIRA regulates transcription and establish HIRA’s critical role in insulin sensitivity and lipid metabolism, providing a potential therapeutic target for obesity and insulin resistance.

**Significance Statement:** Dysregulated adipose tissue function leads to metabolic disorders including obesity and insulin resistance, yet the epigenomic mechanisms that maintain its normal function are incompletely understood. This study identifies the histone chaperone HIRA as a critical regulator of insulin sensitivity and obesity-associated adipose expansion. We show that HIRA directly controls adiponectin and lipid metabolism gene expression in adipocytes by promoting RNA polymerase II pause release, rather than transcription initiation or enhancer activation, and likely in a manner independent of H3.3 deposition. These findings uncover a novel mechanism of HIRA in regulating gene expression and highlight its critical role in insulin sensitivity and lipid metabolism in adipocytes.

## Introduction

Adipose tissue plays an essential role in various physiological processes including energy storage, insulin sensitivity, and glucose and lipid homeostasis in mammals (1, 2). Adipose tissue is also considered an endocrine organ and secretes adipokines. Adiponectin, encoded by the *Adipoq* gene, is an adipokine specifically and abundantly expressed in adipose tissue. Adiponectin plays a critical role in regulating insulin sensitivity, as *Adipoq* knockout (KO) mice exhibit severe insulin resistance, especially when challenged with a high-fat diet (HFD) (3, 4). Development of adipose tissue (adipogenesis) is under the control of transcription factors (TFs) and epigenomic regulators (5). The master adipogenic TF PPARγ (peroxisome proliferator-activated receptor gamma) cooperates with another key TF C/EBPα (CCAAT enhancer-binding protein alpha) on transcriptional enhancers to induce expression of thousands of adipocyte genes (6). Among them, genes encoding lipogenic TFs and enzymes are induced in the late phase of adipogenesis. Increases in adipocyte number due to adipogenesis (hyperplasia), and more importantly enlarged adipocytes (hypertrophy) due to lipogenesis or lipid uptake, lead to adipose tissue expansion in obesity (2, 7).

Adipose tissue and the liver are major lipogenic organs and coordinate the conversion of excess dietary carbohydrates into fatty acids through glycolysis, the tricarboxylic acid cycle, and de novo lipogenesis. Fatty acids are subsequently esterified into triglyceride for storage in lipid droplets (8–10). Glucose and insulin promote lipogenesis gene expression in liver and adipose tissue through activation of lipogenic TFs ChREBP (carbohydrate response element binding protein) and SREBP1 (sterol regulatory element binding protein 1) (11, 12). ChREBP and SREBP1 directly promote expression of genes encoding lipogenesis enzymes ATP-citrate lyase (ACLY), acetyl-CoA carboxylase (ACACA), fatty acid synthase (FASN), and stearoyl-CoA desaturase 1 (SCD1), resulting in fatty acid synthesis and triglyceride accumulation in the late phase of adipogenesis (10). However, the role of histone chaperones in adipose tissue function and expansion has not been reported.

The histone chaperone HIRA (histone regulator A) deposits the histone H3 variant H3.3 into gene regulatory regions including enhancers and promoters in a DNA synthesis-independent manner (13–17). ChIP-Seq analyses in human HeLa and mouse embryonic stem (ES) cells reveal that HIRA colocalizes with H3.3 on promoters and active enhancers (AEs) (14, 18) and knockdown of *HIRA* expression markedly reduces H3.3 deposition at these sites (18, 19). Depletion of HIRA does not affect ES cell self-renewal and only leads to mild changes in global gene expression, but impairs neural differentiation (14, 19). *Hira* KO mouse embryos show abnormal gastrulation and die before embryonic day 11 (20). HIRA is required for normal heart development and generation of all hematopoietic lineages in mice (21, 22). Adult mice with muscle-specific deletion of *Hira* show impaired muscle regeneration (23). These findings suggest a lineage– or stage-selective role of HIRA in differentiation and development.

In this study, we explored the role of HIRA in adipose tissue function and expansion. By crossing *Hira*^f/f^ with *Adipoq-Cre* mice, we observed that deletion of *Hira* in adipocytes significantly reduces serum adiponectin levels, leads to insulin resistance under normal chow diet, and renders mice resistant to adipose tissue expansion during HFD-induced obesity. RNA-Seq and ChIP-Seq analyses revealed that HIRA directly binds to promoters and enhancers of *Adipoq* and lipid metabolism genes to regulate their expression in adipocytes. To understand the underlying molecular mechanism, we employed the dTAG rapid protein degradation system coupled with nascent RNA-Seq and ChIP-Seq in adipocytes. dTAG-mediated acute depletion of HIRA has minimal effects on the binding of coactivators CBP, BRD4 and MED1 on promoters and AEs. Interestingly, HIRA depletion impairs RNA polymerase II (Pol II) pause release and subsequent transcription elongation at target genes likely independently of H3.3 deposition. These findings reveal a previously unrecognized mechanism of HIRA-mediated transcriptional regulation and establish a critical role of HIRA in regulating insulin sensitivity and lipid metabolism in adipocytes.

## Results

### Mice with adipocyte-specific deletion of *Hira* show reduced adiponectin levels and insulin resistance under normal chow diet

To investigate the functional role of HIRA in adipose tissues, we generated adipocyte-specific *Hira* KO (*Hira*^f/f^*;Adipoq-Cre* [A-KO]) mice by crossing *Hira*^f/f^ with *Adipoq-Cre* mice. qRT-PCR confirmed the specific reduction of *Hira* expression in adipose tissues but not liver (Fig. S1a). A-KO and f/f mice showed similar appearance, body weight, fat and lean mass (Fig. 1a-b). The weight and size of adipose tissues including interscapular white adipose tissue (intWAT), brown adipose tissue (BAT), inguinal WAT (ingWAT) and epididymal WAT (eWAT) and liver were similar between f/f and A-KO mice (Fig. 1c-d). Histological analysis also revealed comparable lipid droplets in adipose tissues and liver in f/f and A-KO mice (Fig. S1b).

**Figure 1.**
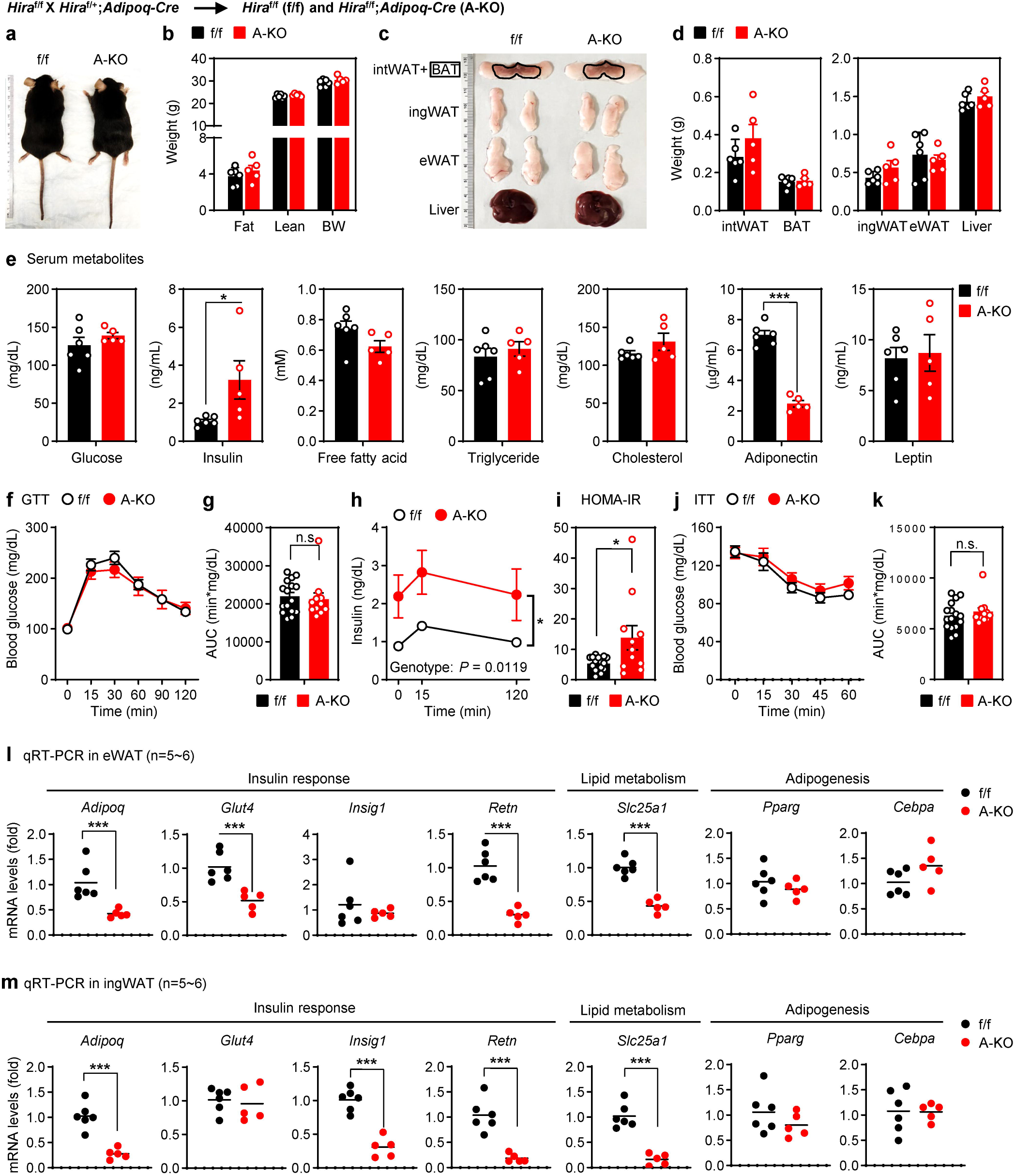
Mice with adipocyte-specific deletion of Hira show reduced adiponectin levels and insulin resistance under normal chow diet. All data were from 23-week-old Hira^f/f^ (f/f) and Hira^f/f^;Adipoq-Cre (A-KO) male mice fed with a normal chow diet (a-e, l-m, *n* = 5∼6 per group; f-k, *n* = 11∼16 per group). **a**, Representative morphology of mice. **b**, Body composition measured by MRI. **c**, Representative pictures of interscapular WAT (intWAT), BAT, inguinal WAT (ingWAT), epididymal WAT (eWAT) and liver. **d**, Average tissue weights. **e**, Levels of serum metabolites in randomly fed state. **f-h**, Glucose tolerance test (GTT): blood glucose levels (**f**), area under the curve (AUC, **g**), and insulin levels (**h**). **i**, Insulin sensitivity was determined by HOMA-IR. **j-k**, Insulin tolerance test (ITT): blood glucose levels (**j**) and area under the curve (AUC, **k**). **l-m**, qRT-PCR analysis of representative genes in eWAT (**l**) and ingWAT (**m**). All quantitative data for mice are presented as means ± SEM. Statistical comparison between groups was performed using Student’s *t*-test (**e**, **g**, **i, k, l** and **m**) and two-way repeated measures ANOVA with Sidak post hoc analysis (**f, h** and **j**). (∗) *P* < 0.05, (∗∗) *P* < 0.01, (∗∗∗) *P* < 0.001.

Interestingly, non-fasting serum insulin levels were significantly elevated while adiponectin levels were around 3-fold lower in male A-KO compared to f/f mice (Fig. 1e). No significant differences were observed in glucose, free fatty acid, triglyceride, cholesterol and leptin levels in A-KO mice. We then performed glucose tolerance test (GTT) and insulin tolerance test (ITT). Fasting blood glucose, glucose excursion curves during GTT, and the reduction of blood glucose during ITT were comparable between A-KO and f/f mice (Fig. 1f-g & j-k). However, A-KO mice exhibited significantly elevated insulin levels during GTT, including fasting insulin at 0 min (Fig. 1h), along with a higher HOMA-IR (Homeostatic Model Assessment for Insulin Resistance) index (Fig. 1i), while expression of glucose uptake genes remain largely unchanged in the liver (Fig. S1c). Taken together, these data are consistent with a modest insulin resistance in male A-KO mice. Female A-KO mice displayed similar metabolic phenotype, including low serum adiponectin levels, elevated insulin during GTT, and impaired insulin tolerance with higher glucose levels during ITT (Fig. S2). Additionally, A-KO and f/f mice showed similar levels of energy expenditure and lipolysis (Fig. S1d-e). Taken together, these data suggest that HIRA is important for glucose homeostasis and adipocyte-specific deletion of *Hira* leads to features consistent with impaired insulin sensitivity.

Next, we performed RNA-Seq analysis in eWAT and ingWAT. Using a 1.5-fold cutoff for differential gene expression, we identified 215 downregulated and 442 upregulated genes in eWAT, and 95 downregulated and 73 upregulated genes in ingWAT of A-KO mice compared to f/f mice (Fig. S3a-b). We found 48 genes downregulated over 1.5-fold in both eWAT and ingWAT of A-KO mice (Fig. S3c & Supplemental Table 1), including genes involved in insulin response and lipid metabolism, such as *Adipoq*, *Retn* and *Slc25a1* (Fig. S3d-e). qRT-PCR further validated decreased expression of *Adipoq*, *Glut4*, *Insig1*, *Retn* and *Slc25a1* in eWAT and/or ingWAT of A-KO mice. In contrast, the expression of adipogenesis markers *Pparg* and *Cebpa* remained unchanged (Fig. 1l-m). Since reduced adiponectin levels impair insulin sensitivity in mice (3, 24), our data suggest that HIRA may regulate insulin response/sensitivity, at least in part, by controlling *Adipoq* expression.

### Mice with adipocyte-specific deletion of *Hira* gain less fat mass during HFD-induced obesity

Next, we examined the role of HIRA in diet-induced obesity. Eight-week-old f/f and A-KO mice were fed with HFD for 8 weeks. qRT-PCR confirmed the reduction of *Hira* expression in adipose tissues but not liver after 8 weeks on the HFD (Fig. S4a). A-KO mice gained a similar amount of lean mass, but significantly less fat mass and body weight compared to f/f mice (Fig. 2a-d) with lower cumulative food intake starting at week 4 and largely unchanged energy expenditure (Fig. S4b-c). Among adipose tissues examined, A-KO mice showed less gain of eWAT compared to f/f mice (Fig. 2e-f). Histological analysis revealed a substantially higher proportion of smaller adipocytes in eWAT and ingWAT of A-KO mice (Fig. 2g-h). Under HFD condition, A-KO and fl/fl mice exhibited comparable levels of glucose, insulin, and HOMA-IR, as well as similar responses to exogenous glucose (in GTT) and insulin (in ITT) (Fig. 2i & Fig. S4d-i). However, serum levels of total cholesterol, adiponectin and leptin were significantly lower in A-KO mice (Fig. 2i).

**Figure 2.**
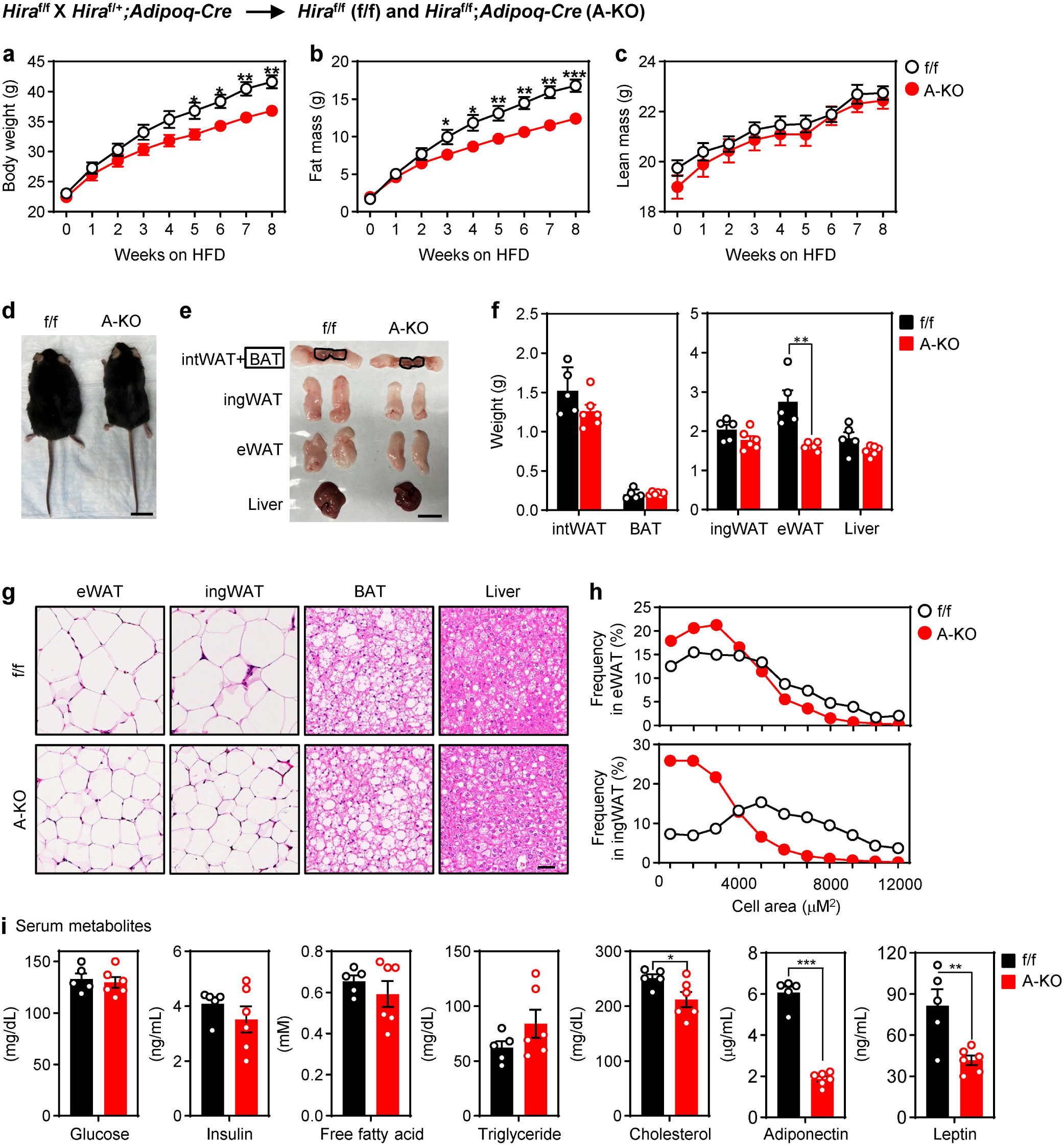
Mice with adipocyte-specific deletion of *Hira* gain less fat mass during HFD-induced obesity. Male *Hira*^f/f^ (f/f) and *Hira*^f/f^;*Adipoq-Cre* (A-KO) mice (*n* = 5∼6 per group) were fed with HFD from the eighth week of age. **a-c**, Total body weight (**a**), fat mass (**b**), and lean mass (**c**) were measured by MRI during HFD feeding. **d**, Representative morphology of HFD-fed mice. Scale, 2 cm. **e**, Representative pictures of intWAT, BAT, ingWAT, eWAT and liver. Scale, 2 cm. **f**, Average tissue weights. **g**, H&E staining of eWAT, ingWAT, BAT and liver. Scale bar, 50 μm. **h**, Average cell area in eWAT (upper) and ingWAT (lower). **i**, Levels of serum metabolites. All quantitative data for mice are presented as means ± SEM. Statistical comparison between groups was performed using Student’s *t*-test. (∗) *P* < 0.05, (∗∗) *P* < 0.01, (∗∗∗) *P* < 0.001.

We then performed RNA-Seq in eWAT and ingWAT after HFD. A-KO mice showed the loss of RNA-Seq signals on the floxed exon 7 of *Hira*, indicating successful deletion (Fig. 3a). Using a 1.5-fold cutoff for differential gene expression, we identified 374 downregulated and 258 upregulated genes in eWAT, and 505 downregulated and 352 upregulated genes in ingWAT of A-KO mice compared to f/f mice (Fig. 3b-c). Among these, 175 genes were consistently downregulated by more than 1.5-fold in both eWAT and ingWAT of A-KO mice (Fig. 3d), showing functional enrichment in lipid metabolic process (Fig. 3e & Fig. S5a). In addition to *Adipoq*, expression levels of lipid metabolism genes, such as *Fasn*, *Agpat2*, *Bscl2*, *Cd36* and *Fatp1* were reduced in both eWAT and ingWAT of A-KO mice (Fig. 3f-g & Fig. S5b). These results suggest that HIRA plays an important role in adipose tissue expansion during HFD-induced obesity, at least in part, by regulating *Adipoq* and lipid metabolism genes.

**Figure 3.**
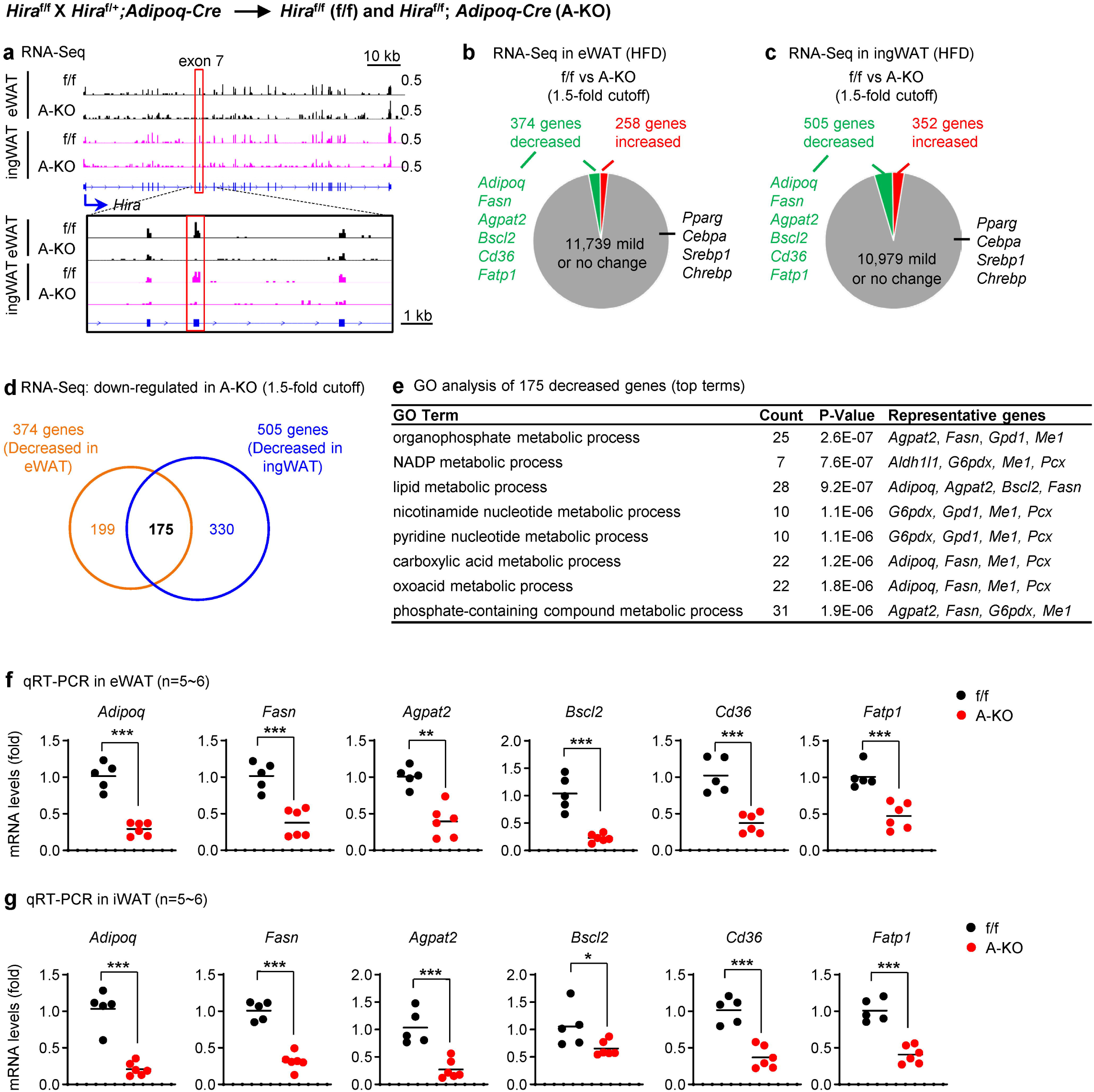
Adipocyte-specific deletion of *Hira* reduces expression of *Adipoq* and lipid metabolism genes in WAT during HFD-induced obesity. All data were from *Hira*^f/f^ (f/f) and *Hira*^f/f^;*Adipoq-Cre* (A-KO) mice (*n* = 5∼6 per group) fed with HFD for 8 weeks from the eighth week of age. **a**, Genome browser view of RNA-Seq data on the *Hira* gene locus. The floxed exon 7 is highlighted. **b-c**. RNA-Seq analysis of eWAT (**b**) and ingWAT (**c**). Differentially expressed genes were identified using DESeq2 (35) in R (v3.5.3), applying a threshold of 1.5-fold change and a *P*-value < 0.01. **d**, Venn diagrams depicting down-regulated genes in adipose tissues of A-KO mice. **e**, GO analysis of gene groups defined in **d**. **f-g**, qRT-PCR analysis of representative genes in eWAT (**f**) and ingWAT (**g**). All quantitative data for mice are presented as means ± SEM. Statistical comparison between groups was performed using Student’s *t*-test. (∗) *P* < 0.05, (∗∗) *P* < 0.01, (∗∗∗) *P* < 0.001. RNA-Seq analysis was done using two independent samples (biological replicates) for each genotype. Each sample was a pool of RNA isolated from two to three mice.

### Knockout of *Hira* in white preadipocytes prevents adipogenesis

To understand how HIRA regulates expression of insulin response genes including *Adipoq* and lipid metabolism genes, we generated *Hira* KO cells by CRISPR/Cas9 in 3T3-L1 white preadipocytes using lentiviral *Hira* gRNA (Fig. S6a). KO of *Hira* in 3T3-L1 led to adipogenesis defects as shown by severely reduced Oil red O staining (Fig. S6b). RNA-Seq analysis showed that 1,710 (15.2%) and 1,755 (15.6%) genes were over 2-fold up– and down-regulated from day 0 (D0) to day 7 (D7), respectively, in the control cells. Among the 1,710 up-regulated genes, 1,280 (74.9%) were induced in a HIRA-dependent manner (Fig. S6c). Consistent with *in vivo* data, Gene Ontology (GO) analysis showed that the term “lipid metabolic process” was top-ranked in the HIRA-dependently up-regulated gene group (Fig. S6d). Notably, the induction of adipogenesis markers *Pparg* and *Cebpa* as well as insulin response gene *Adipoq* and the lipogenesis gene *Fasn* was prevented in *Hira* KO cells at D7 (Fig. S6e). These data suggest that HIRA is required for adipogenesis in culture.

Because the impaired induction of insulin response and lipid metabolism genes could be secondary to the adipogenesis defect in culture, we employed a dTAG-13–mediated protein degradation system in 3T3-L1 cells to allow temporal control of HIRA depletion. Exogenous HIRA protein tagged with C-terminal dTAG and HA epitopes (HIRA-dTAG-HA) was introduced into 3T3-L1 cells, followed by CRISPR/Cas9-mediated deletion of the endogenous *Hira* gene (Fig. S6f). Adipogenesis assays demonstrated that HIRA-dTAG-HA could functionally substitute for endogenous HIRA (Fig. S6g–h).

### HIRA directly binds insulin response and lipid metabolism genes in adipocytes

The dTAG-13 mediated protein degradation approach enabled us to deplete HIRA during adipogenesis (Fig. 4a) (25) and map high-confidence genomic binding regions of HIRA using a ChIP-Seq quality HA antibody. Western blotting and ChIP-Seq confirmed efficient depletion of HIRA-dTAG-HA before (day 0, D0), during (day 4, D4), and after (day 7, D7) adipogenesis (Fig. 4b-c). ChIP-Seq analysis successfully identified 63,401, 28,449, and 34,591 HIRA binding regions at D0, D4 and D7 of adipogenesis, respectively. dTAG-13 treatment for 24 h eliminated over 90% of HIRA-HA genomic binding. At D0, D4 and D7, 63,375 out of 63,401 (99.96%), 28,021 out of 28,449 (98.50%) and 32,400 out of 34,591 (93.67%) HIRA-binding regions exhibited a more than 2-fold decrease in HIRA binding, respectively (Fig. 4c). Unbiased K-means clustering analysis revealed that genes with significantly increased HIRA binding at D4 were functionally associated with “cellular response to insulin stimulus” and “fatty acid metabolic process” (Fig. 4d & Fig. S7a). Genomic tracks showed that HIRA directly bound to genes encoding adiponectin, ChREBP and FASN at D4 (Fig. 4e), a time point at which *Adipoq* and *Chrebp* were already strongly induced (Fig. 4f). Adipogenesis markers *Pparg* and *Cebpa* were induced at D2, while *Hira* levels remained relatively unchanged during adipogenesis (Fig. 4f). Together, these results indicate that HIRA directly targets insulin response and lipid metabolism genes in differentiating adipocytes at D4. These data also suggest that depleting HIRA at D4 can bypass early adipogenesis defects, allowing the investigation of its role in adipocytes.

**Figure 4.**
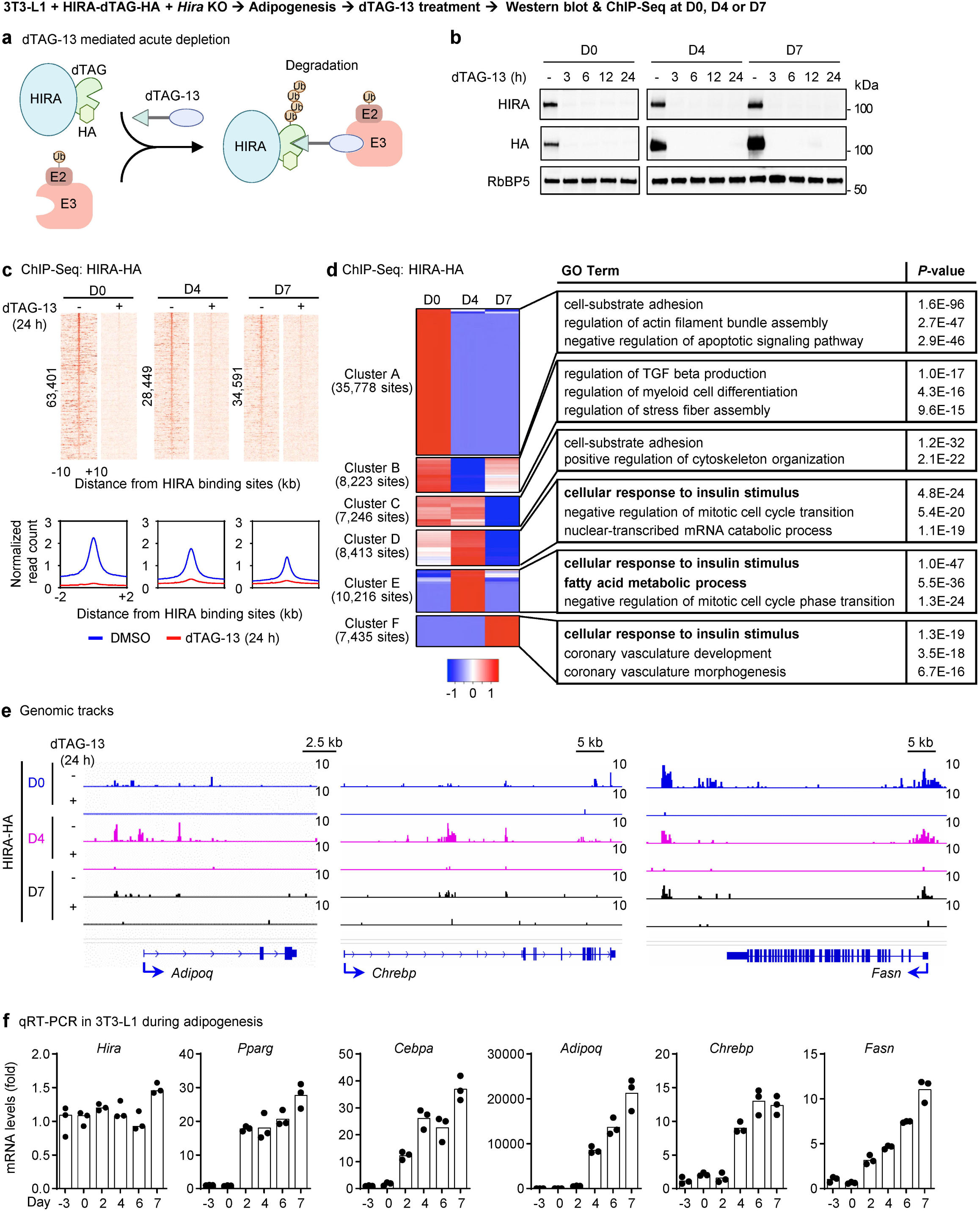
HIRA directly binds insulin response and lipid metabolism genes in adipocytes. 3T3-L1 white preadipocytes were infected with a lentiviral vector expressing HIRA with C-terminal dTAG and HA double tags, followed by lentiviral CRISPR/Cas9-*Hira* gRNA infection to delete endogenous *Hira*. Adipogenesis assay was performed, followed by dTAG-13 treatment at D0, D4, or D7. **a,** A schematic illustration of dTAG-13 mediated acute depletion of HIRA. The chemical dTAG-13 binds to HIRA and recognizes E3 ligase complex, leading to HIRA ubiquitination by an E2 conjugating enzyme and subsequent degradation by the proteasome. **b**, Western blot analysis using indicated antibodies. Cells were treated with dTAG-13 for various time periods as indicated. RbBP5 was used as a loading control. **c-e,** ChIP-Seq analysis of HIRA genomic binding in the presence or absence of dTAG-13 treatment for 24 h. **c**, Heat maps (upper) and average profiles (lower) were aligned around the center of HIRA binding sites at D0 (63,401), D4 (28,449) and D7 (34,591). **d**, Heatmaps of K-means clustering depicting HIRA binding regions at D0, D4 and D7. **e**, ChIP-Seq profiles of HIRA binding on *Adipoq*, *Chrebp* and *Fasn* gene loci at D0, D4 and D7. **f**, Time course qRT-PCR analysis of representative genes during adipogenesis (*n* = 3).

### Acute depletion of HIRA reduces transcription of *Adipoq* and lipid metabolism genes in adipocytes

To investigate the direct role of HIRA in regulating target gene expression, we depleted HIRA at D4 using dTAG-13 treatment for 6 or 24 h and collected newly synthesized RNA for nascent RNA sequencing (nascent RNA-Seq) (Fig. 5a). We focused on genes that were both upregulated at D4 and directly targeted by HIRA. To this end, we first identified 1,601 genes that exhibited more than a two-fold increase in expression from D0 to D4 during 3T3-L1 differentiation, as determined by RNA-Seq analysis (Fig. S8a). By overlapping these induced genes with the set of HIRA target genes at D4 (Fig. 4d), we identified 1,027 genes that were not only upregulated at D4 but also directly bound by HIRA (Fig. S8b–d). We classified these 1,027 genes into four groups (Q1 – Q4) based on their changes in transcription following HIRA depletion (Fig. 5b). Interestingly, genes in the Q1 group, which exhibited the most significant downregulation after 6-hour and 24-hour HIRA depletion, were strongly associated with lipid metabolic process (Fig. 5c & Fig. S9a-f). Notably, transcription of *Adipoq*, lipid metabolism genes including *Agpat2*, *Bscl2*, *Dgat1*, *Gpd1,* and lipogenic TF *Chrebp* and *Srebp1* was downregulated following 6-hour and 24-hour HIRA depletion (Fig. 5d & Fig. S9g-h). These data indicate that HIRA directly promotes *Adipoq* and lipid metabolism gene transcription in differentiating adipocytes.

**Figure 5.**
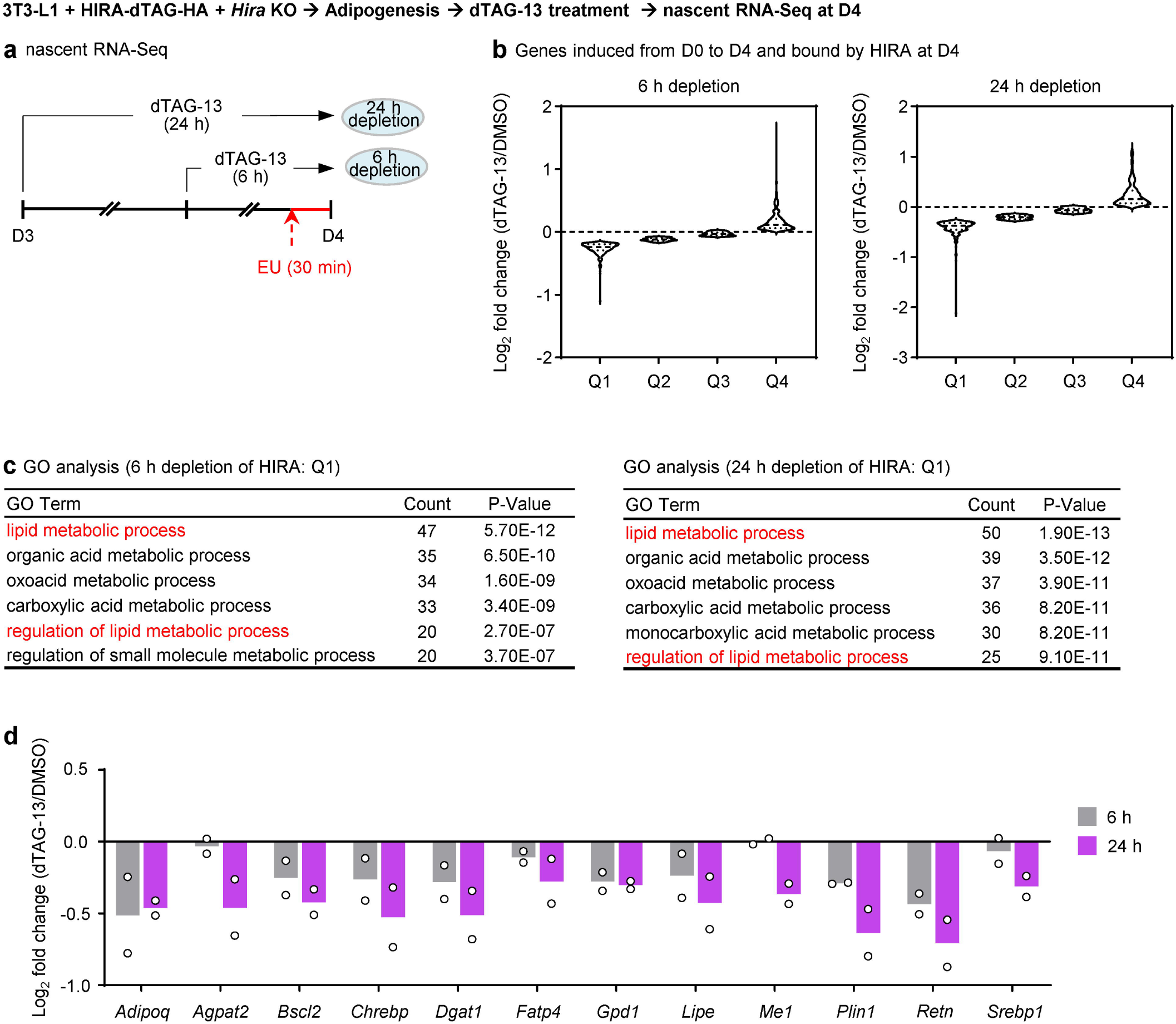
Acute depletion of HIRA reduces transcription of *Adipoq* and lipid metabolism genes in adipocytes. 3T3-L1 white preadipocytes were infected with a lentiviral vector expressing HIRA with C-terminal dTAG and HA double tags, followed by lentiviral CRISPR/Cas9-*Hira* gRNA infection to delete endogenous *Hira*. dTAG-13 treatment was done at D4, followed by nascent RNA-Seq (ncRNA-Seq) analysis. **a,** Schematic of ncRNA-Seq analysis following HIRA depletion for 6 h or 24 h. **b**, Violin plots showing transcriptional changes of the 1,027 genes identified in Fig. S8b following 6 h (left) or 24 h (right) depletion of HIRA (*n* = 2). Genes were ranked according to their transcriptional changes upon HIRA depletion and divided into four quartiles (Q1-Q4), ranging from the most downregulated to the most upregulated. **c**, GO analysis of genes in Q1 following 6 h (left) and 24 h (right) depletion of HIRA. **d**, Bar graph showing representative gene transcription changes upon depletion of HIRA.

### Acute depletion of HIRA does not interfere with the recruitment of transcription coactivators

To understand how HIRA regulates transcription, we further characterized the genomic distribution of HIRA in adipocytes. By ChIP-Seq analysis of enhancer mark H3K4me1, active enhancer (AE) mark H3K27ac, and HIRA-HA at D4 of adipogenesis, we found that 6,838 out of 28,499 HIRA binding sites were on promoters, encompassing +/− 1 kb relative to the transcription start site (TSS). 17,862 (62.8%) sites were on AEs, which are promoter-distal regions marked by H3K4me1 and H3K27ac (Fig. 6a). To minimize potential secondary effects caused by permanent *Hira* gene deletion, we utilized 6-hour dTAG-13 treatment to acutely deplete HIRA protein at the genomic level at D4 of adipogenesis. Interestingly, despite HIRA’s predominant binding to these regulatory elements, its acute depletion had no impact on H3K27ac enrichment—a hallmark of active regulatory elements —or the binding of its cognate acetyltransferase CBP, suggesting that HIRA is dispensable for maintaining AEs (Fig. 6b-c). We next examined the impact of HIRA depletion on downstream coactivators, including BRD4 and the Mediator complex subunit MED1 (10, 26). Although a modest increase in BRD4 and MED1 occupancy was observed at HIRA^+^ promoters, their binding at HIRA^+^ AEs remained unchanged upon acute HIRA depletion (Fig. 6b–c). Consistently, H3K27ac, CBP, BRD4, and MED1 occupancy at representative metabolic gene loci, including insulin response gene *Adipoq*, lipogenic TF *Chrebp* and lipogenesis gene *Fasn*, was unaffected by acute HIRA loss (Fig. 6d). Taken together, our data indicate that acute depletion of HIRA does not interfere with the recruitment of transcription coactivators.

**Figure 6.**
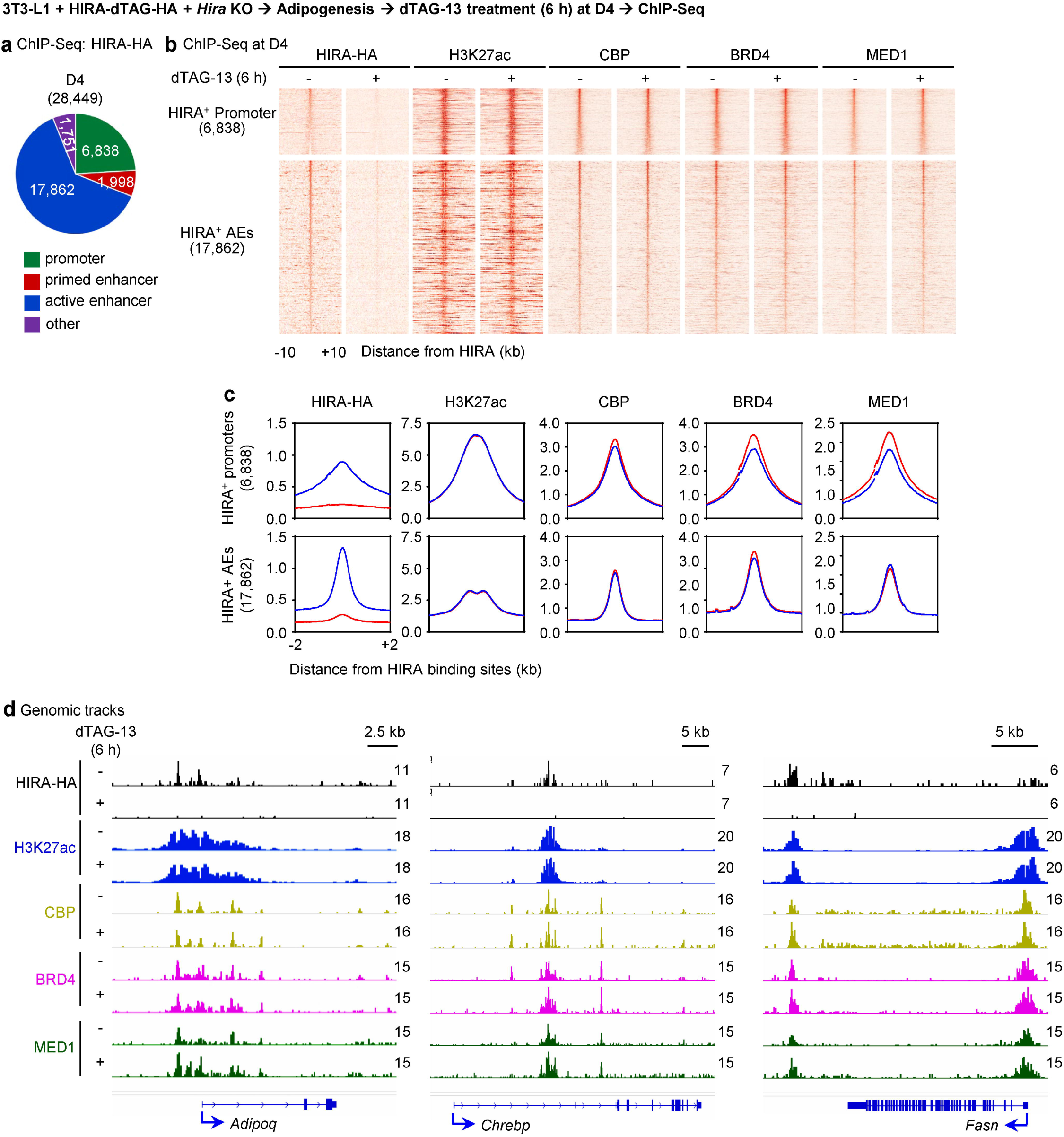
Acute depletion of HIRA does not interfere with the recruitment of transcription coactivators. 3T3-L1 white preadipocytes were infected with a lentiviral vector expressing HIRA with C-terminal dTAG and HA double tags, followed by lentiviral CRISPR/Cas9-*Hira* gRNA to delete endogenous *Hira*. At D4 of adipogenesis, cells were treated with dTAG-13 for 6 h. **a**, Pie charts depicting the genomic distribution of HIRA binding regions at D4. **b–c,** Heat maps (**b**) and average profiles (**c**) showing HIRA-HA binding, H3K27ac enrichment, and CBP, BRD4 and MED1 occupancy aligned to the centers of HIRA binding sites at promoters (6,838) and active enhancers (AEs; 17,862) at D4. **d**, ChIP–Seq profiles of HIRA-HA, H3K27ac, CBP, BRD4 and MED1 were displayed on *Adipoq*, *Chrebp* and *Fasn* loci.

### Acute depletion of HIRA impairs RNA Pol II pause release on insulin response and lipid metabolism genes in adipocytes

Next, we examined RNA polymerase II (Pol II) occupancy upon 6 h depletion of HIRA (Fig. S10a). We observed a significant increase of Pol II binding on HIRA^+^ promoters, with a globally increased pausing index (Fig. 7a-b), suggesting a defect in Pol II promoter pause release upon HIRA loss. In accordance, enrichment of initiating Pol II, indicated by serine-5 phosphorylation (S5P), increased on HIRA^+^ promoters, AEs and transcribed gene bodies, while enrichment of elongating Pol II, defined by serine-2 phosphorylation (S2P), decreased after HIRA depletion for 6 h (Fig. 7c). Similarly, transcription elongation factors CDK9 and SPT6 decreased at enhancers/promoters and throughout the gene bodies upon HIRA loss. Binding of promoter-proximal pausing factor NELF on HIRA^+^ promoters and AEs remained unchanged (Fig. 7c & Fig. S10b). Although HIRA protein was almost completely depleted within 3 h (Fig. 4b), substantial levels of H3.3 on HIRA^+^ sites including promoters persisted after 6 h of HIRA depletion but exhibited a pronounced reduction after 48 h of depletion (Fig. 7d & Fig. S10c).

**Figure 7.**
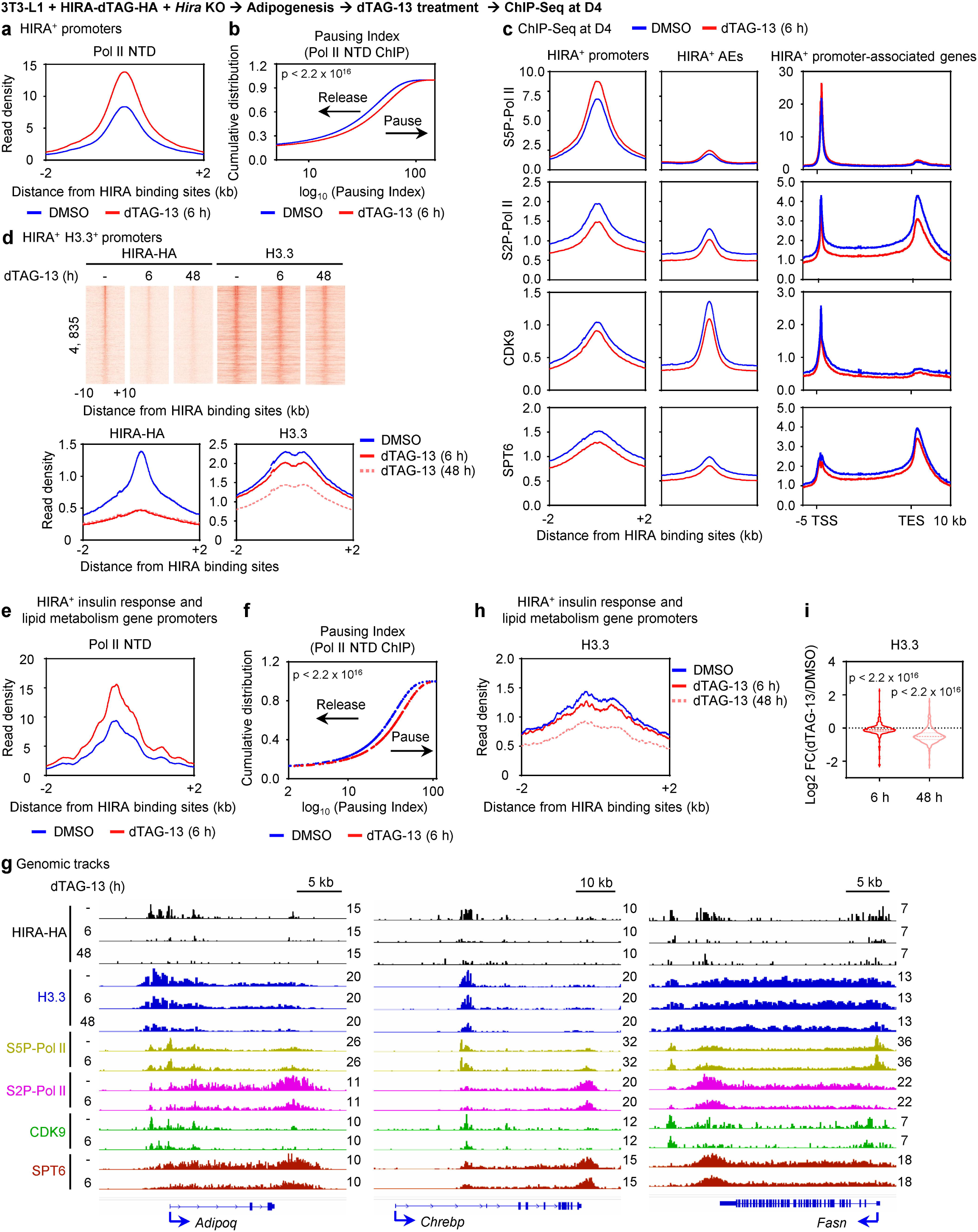
Acute depletion of HIRA impairs RNA Pol II pause release on insulin response and lipid metabolism genes in adipocytes. 3T3-L1 white preadipocytes were infected with a lentiviral vector expressing HIRA with C-terminal dTAG and HA double tags, followed by lentiviral CRISPR/Cas9-*Hira* gRNA to delete endogenous *Hira*. dTAG-13 treatment was done at D4, followed by ChIP-Seq analysis. **a**, Average binding profile of total Pol II (Pol II NTD) around the center of HIRA^+^ promoters after 6 h depletion of HIRA. **b**, Cumulative distribution graph of Pol II pausing index following HIRA depletion for 6 h. *P-*value was calculated using paired Wilcoxon signed-rank test. **c**, Average binding profiles of S5P-Pol II, S2P-Pol II, CDK9 and SPT6 on HIRA^+^ promoters, AEs and HIRA^+^ promoter-associated gene bodies. **d,** Heat maps (upper) and average profiles (lower) showing HIRA-HA binding and H3.3 enrichment aligned to the center of HIRA^+^ H3.3^+^ promoters. **e-i,** ChIP-Seq analysis on HIRA^+^ insulin response and lipid metabolism genes identified in Fig. S7a. **e**, Average binding profile of total Pol II (Pol II NTD) aligned to the center of HIRA^+^ promoters. **f**, Cumulative distribution graph of Pol II pausing index. *P-*value was calculated using paired Wilcoxon signed-rank test. **g**, ChIP–Seq profiles of HIRA-HA, H3.3, S5P-Pol II, S2P-Pol II, CDK9 and SPT6 on *Adipoq*, *Chrebp* and *Fasn* loci. **h**, Average binding profile of H3.3 aligned to the center of HIRA^+^ promoters. **i**, Quantification of (**h**) is presented as violin plots showing the fold changes of H3.3 intensities on TSS ± 2 kb. *P*-value was calculated using paired Wilcoxon signed-rank test.

We next focused on HIRA^+^ insulin response and lipid metabolism genes as identified in Fig. S7a. Consistent with global trends observed for all HIRA^+^ promoters, we detected a significant increase in promoter Pol II occupancy, accompanied by an elevated pausing index (Fig. 7e–f). Consistently, decreased levels of S2P-Pol II, CDK9, and SPT6 were observed on enhancers and promoters of *Adipoq*, *Chrebp* and *Fasn* (Fig. 7g). Similar to the pattern observed at HIRA⁺ H3.3^+^ promoters, H3.3 deposition on promoters of insulin response and lipid metabolism genes exhibited only a mild reduction following 6 h of HIRA depletion but a pronounced decrease after 48 h of depletion (Fig. 7h-i). Although we cannot exclude the possibility that subtle reductions in H3.3 following 6 h depletion of HIRA contribute to Pol II pause release defects, our data suggest that HIRA regulation of Pol II pause release on insulin response and lipid metabolism genes is largely independent of H3.3 deposition.

## Discussion

In this study, using adipose tissue-specific gene KO in mice, we identify the histone chaperone HIRA as a novel epigenomic regulator of insulin sensitivity and obesity-associated adipose tissue expansion. Deletion of *Hira* reduces expression of insulin sensitivity gene *Adipoq* and lipid metabolism genes in adipocytes in mice. By ChIP-Seq, we determine genomic binding sites of HIRA and show that HIRA targets promoters and enhancers of *Adipoq* and lipid metabolism genes in adipocytes. Finally, acute HIRA protein depletion in adipocytes reveals that HIRA is dispensable for coactivators’ binding, but required for Pol II pause release and subsequent transcription elongation, on target genes, providing a novel mechanism by which HIRA regulates transcription. While a previous study suggests that HIRA mediated H3.3 deposition is implicated in Pol II mediated transcription (27), we propose a model that HIRA facilitates the transcription of *Adipoq* and lipid metabolism genes by promoting RNA Pol II pause release and elongation, likely in a H3.3-independent manner (Figure 8). This study reveals a novel transcriptional mechanism regulating lipid metabolism genes, highlighting an unrecognized layer of epigenomic control and providing a potential therapeutic target for obesity.

**Figure 8.**
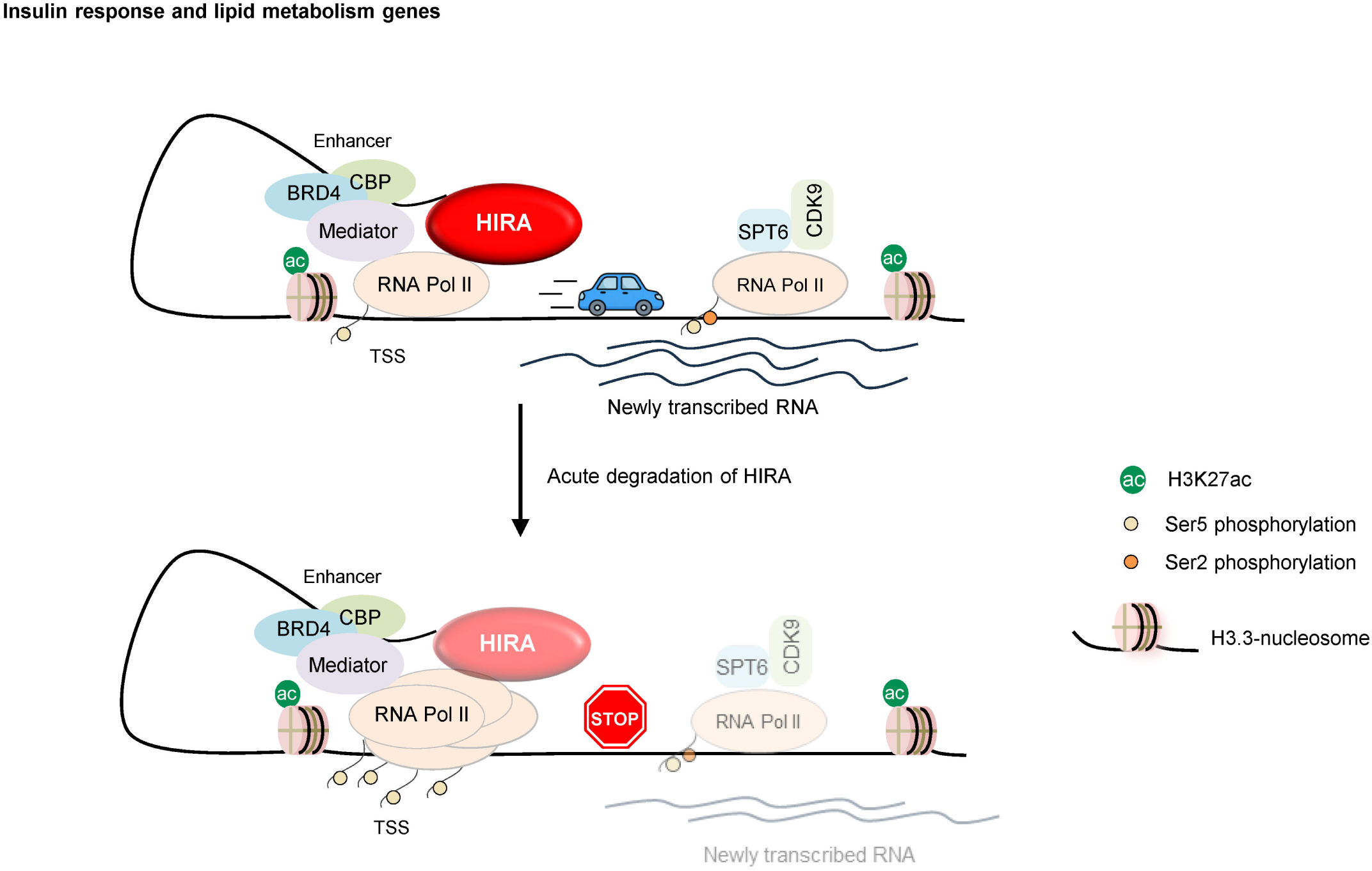
Proposed model. In the presence of HIRA, RNA Pol II efficiently undergoes pause release and enters productive elongation, resulting in efficient RNA synthesis. Upon acute HIRA depletion, transcription coactivator bindings are largely unaffected, but RNA Pol II fails to properly pause release, accumulates at promoter regions, and exhibits impaired elongation with reduced RNA transcription, whereas H3.3 levels remain largely unchanged on HIRA^+^ insulin response and lipid metabolism genes.

By crossing *Hira*^f/f^ with *Adipoq-Cre* mice, we observed that under normal chow diet, both male and female A-KO mice maintain normal body and adipose tissue weights but show significantly decreased serum levels of the insulin-sensitizing adipokine adiponectin, which may contribute to the observed insulin resistance phenotype. Reduced expression of lipogenesis genes such as *Slc25a1* in these mice may also contribute to insulin resistance, because insulin resistance is associated with reduced lipogenesis enzyme expression in WAT (8). Under HFD, the reduced fat mass gain and smaller adipocytes observed in A-KO mice can be partially explained by the decreased expression of lipogenesis genes such as *Fasn*. Consistently, it has been reported that adipocyte-specific *Fasn* KO mice maintain normal body weight under normal chow diet but display reduced fat mass and smaller eWAT under HFD (28). Besides this, the reduced cumulative food intake likely contributes to the decreased fat mass and body weight in A-KO mice, which could also be related to low adiponectin levels since adiponectin-deficient mice showed decreased food intake and exhibited resistance to HFD-induced obesity (29). Under HFD, despite decreased adiponectin levels, *Hira* KO mice did not show more severe insulin resistance compared to controls. This could be due to the concurrent approximately 50% reduced leptin levels observed in *Hira* KO mice (Fig. 2i), since a 40%–50% reduction in circulating leptin levels has been reported to enhance insulin sensitivity in mice under HFD (30).

Using adipogenesis as a model system, we examined HIRA genomic localization during cell differentiation. We used a ChIP-Seq quality HA antibody to detect the exogenous HA-tagged HIRA because the available HIRA antibody worked for western blot but failed in ChIP-Seq. In differentiating adipocytes (D4), HIRA predominantly binds to promoters, primed enhancers, and AEs, consistent with previous findings from HeLa cells (18). Motif enrichment analysis of HIRA-bound AEs revealed a predominance of binding sites for TFs associated with active chromatin, including AP-1, C/EBP, NF1, TEAD, and PPAR (Fig. S7b), suggesting that HIRA recruitment may be broadly mediated through interactions with these enhancer-associated factors. HIRA binding sites are highly associated with insulin response and lipid metabolism genes. Notably, HIRA-bound AEs linked to these genes showed a similar but not identical motif landscape compared to all HIRA binding AEs, with enrichment of C/EBP, AP-1, NF1, TEAD, as well as EBF and GR motifs (Fig. S7c). EBF (31) and GR (32) are transcription factors known to be involved in lipid metabolism, suggesting that transcription factors related with lipid metabolism may contribute to HIRA targeting in a context-dependent manner. Although lipogenic TFs such as SREBP1 and ChREBP were not observed in the motif analysis, we cannot rule out the possibility of these TFs’ roles in recruiting HIRA. Since adipogenic and/or lipogenic TFs PPARγ, C/EBPα, SREBP1 and ChREBP directly bind insulin response gene *Adipoq* (Fig. S7d) and lipogenesis genes and regulate their expression (10, 33), future studies will be needed to find out whether any of these TFs are responsible for recruiting HIRA to these target genes. By acute depletion of HIRA, we demonstrated the importance of HIRA in Pol II pause release, rather than initiation, on insulin response gene *Adipoq* and lipid metabolism genes. Future studies investigating the potential interplay between HIRA and pause release factors may provide deeper insights into the mechanism by which HIRA regulates Pol II pause release on these genes.

HIRA mediated Pol II pause release appears largely independent of H3.3 deposition, as we observed only a mild decrease in H3.3 levels on lipid metabolism genes upon HIRA depletion for 6 h (Fig. 7g-i). However, by examining all expressed genes, we identified several genes that exhibit a more rapid loss of H3.3, with reductions at 6 h comparable to those observed at 48 h after HIRA depletion (Fig. S10d). Notably, these genes also show a concomitant decrease in gene expression, suggesting an H3.3-dependent regulatory mechanism. Together, these findings suggest that HIRA can function through both H3.3-dependent and –independent mechanisms to regulate gene expression. Future studies involving acute H3.3 depletion or disruption of the HIRA–H3.3 interaction may further clarify the mechanisms underlying these distinct modes of regulation.

## Materials and Methods

Lentiviral plasmid pLV-EF1a-IRES-hygro–HIRA-dTAG-HA was generated in this study. Cell culture and adipogenesis assay were performed as described (34). Metabolic studies, western blot analysis, qRT-PCR, ChIP-Seq, RNA-Seq, nascent RNA-Seq and computational analysis were done as described (10). All mouse experiments were performed in accordance with the NIH Guide for the Care and Use of Laboratory Animals and approved by the Animal Care and Use Committee of the National Institute of Diabetes and Digestive and Kidney Diseases, NIH. All materials used and detailed experimental procedures can be found in Supplemental Information Materials and Methods.

## Author Contributions

D.W. and K.G. conceived the study. D.W., Y.-K.P., J.-E.L., and O.G. performed the methodology. D.W., J.-E.L., Y.-K.P., S.M., C.A., O.G., and K.G. performed the investigation. J.-E.L., D.W. and G.X. were responsible for the software and performed the formal analysis and data curation. K.O. provided novel reagents. D.W. and K.G. wrote the original draft of the manuscript. D.W., J.-E.L., G.X., and K.G. reviewed and edited the manuscript. K.G. administered the project and acquired the funding.

## Classification

Biological Sciences/Cell Biology

## Data Availability

All study data are included in the article and/or Supplemental Information Materials and Methods.

## Acknowledgements

We thank NIDDK Mouse Metabolism Core for assistance with metabolic phenotyping, the National Heart, Lung, and Blood Institute DNA Sequencing and Genomics Core and the National Institute of Arthritis and Musculoskeletal and Skin Diseases Genomic Technology Section for next-generation sequencing. This work was supported by the Intramural Research Program of NIDDK, National Institutes of Health to K.G. and O.G.

## NIH Acknowledgement Disclaimer Statement

This research was supported by the Intramural Research Program of the National Institute of Diabetes and Digestive and Kidney Diseases (NIDDK) within the National Institutes of Health (NIH). The contributions of the NIH authors were made as part of their official duties as NIH federal employees, are in compliance with agency policy requirements, and are considered Works of the United States Government. However, the findings and conclusions presented in this paper are those of the authors and do not necessarily reflect the views of the NIH or the U.S. Department of Health and Human Services.

