## Supplemental data for "Histone chaperone HIRA regulates adiponectin expression and obesity-associated adipose expansion by facilitating Pol II pause release"

*for*

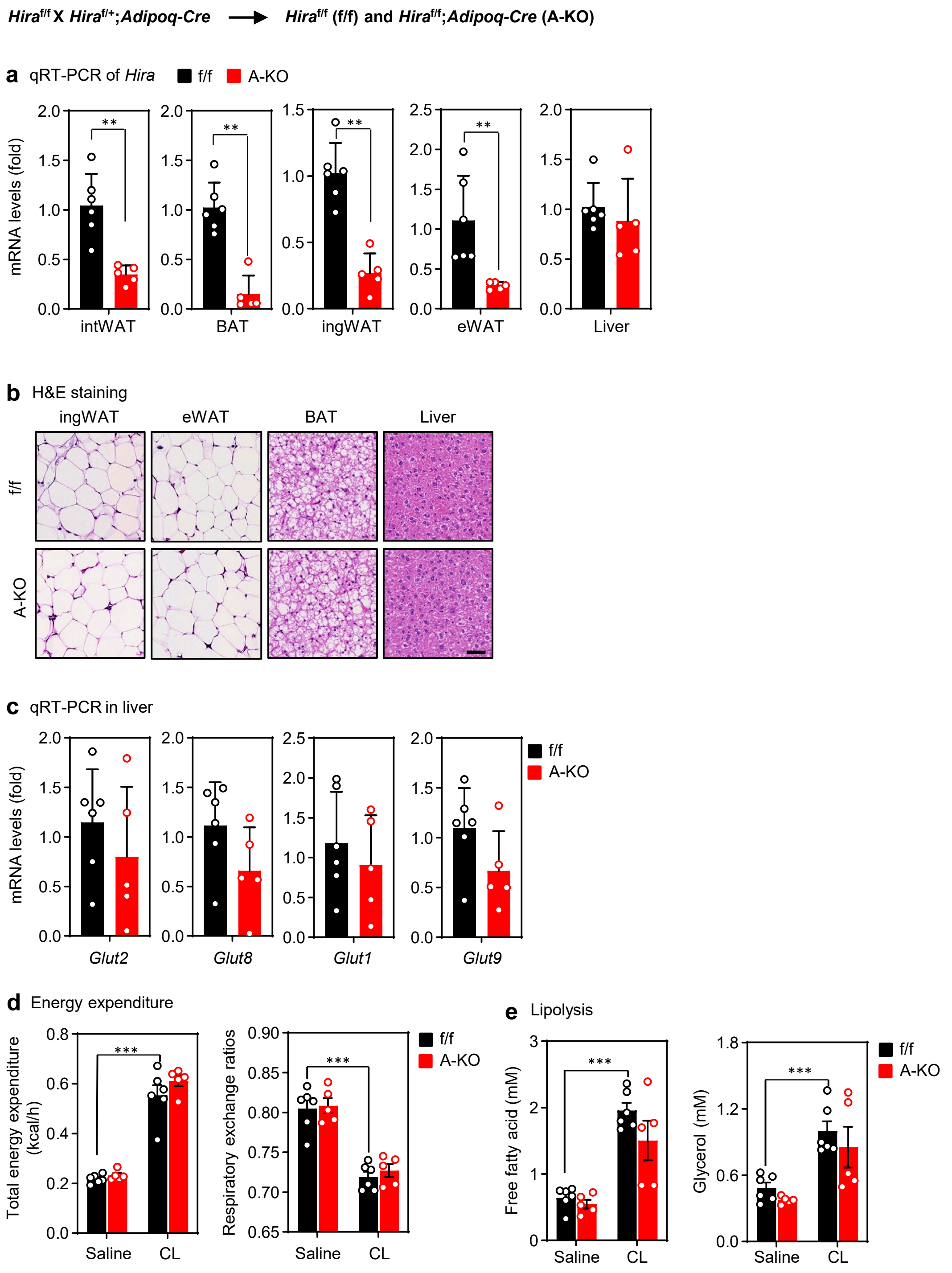


**Figure S1. Characterization of mice with adipocyte-specific deletion of *Hira* under normal chow diet**

All data were from 23-week-old *Hira*$^{f/f}$ (f/f) and *Hira*$^{f/f}$;*Adipoq-Cre* (A-KO) male mice fed with normal chow diet ($n$ = 5~6 per group). **a**, qRT-PCR analysis of *Hira* mRNA levels. **b**, H&E staining of ingWAT, eWAT, BAT and liver. Scale bar, 50 μm. **c**, qRT-PCR analysis of glucose uptake genes in the liver. **d**, Total energy expenditure and respiratory exchange ratios after saline or CL316, 243 (CL) administration. **e**, Lipolysis analysis. Serum levels of free fatty acid or glycerol were measured after saline or CL administration. All quantitative data for mice are presented as means ± SEM. Statistical comparison between groups was performed using Student's $t$-test. (∗) $P < 0.05$, (∗∗) $P < 0.01$, (∗∗∗) $P < 0.001$.

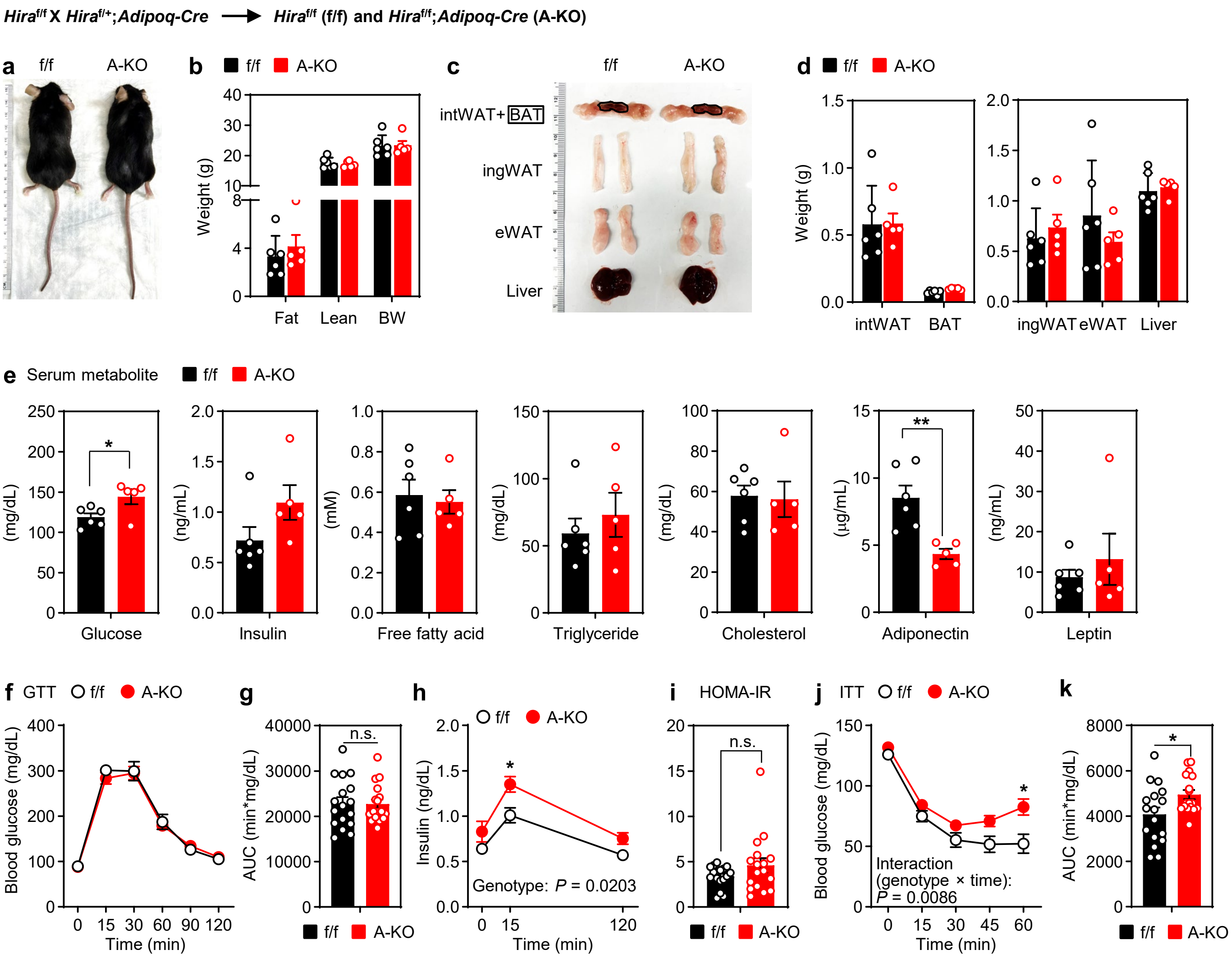


**Figure S2. Female mice with adipocyte-specific deletion of *Hira* show reduced adiponectin levels and insulin resistance under normal chow diet**

All data were from 21-week-old *Hira*$^{f/f}$ (f/f) and *Hira*$^{f/f}$;*Adipoq-Cre* (A-KO) female mice fed with a normal chow diet (**a-e**, *n* = 5~6 per group. **f-k**, *n* = 17 per group). **a**, Representative morphology of mice. **b**, Body composition measured by MRI. **c**, Representative pictures of intWAT, BAT, ingWAT, eWAT and liver. **d**, Average tissue weights. **e**, Levels of serum metabolites. **f-h**, Glucose tolerance test (GTT): blood glucose levels (**f**), area under the curve (AUC, **g**) and insulin levels (**h**). **i**, Insulin sensitivity was determined by HOMA-IR. **j-k**, Insulin tolerance test (ITT): blood glucose levels (**j**) and area under the curve (AUC, **k**). All quantitative data for mice are presented as means ± SEM. Statistical comparison between groups was performed using Student's *t*-test (**e, g, i** and **k**) or two-way repeated measures ANOVA with Sidak post hoc analysis (**f, h** and **j**). (∗) $P < 0.05$, (∗∗) $P < 0.01$, (∗∗∗) $P < 0.001$.

***Hira$^{f/f}$* X *Hira$^{f/+}$;Adipoq-Cre*** ⟶ ***Hira$^{f/f}$* (f/f) and *Hira$^{f/f}$;Adipoq-Cre* (A-KO)**

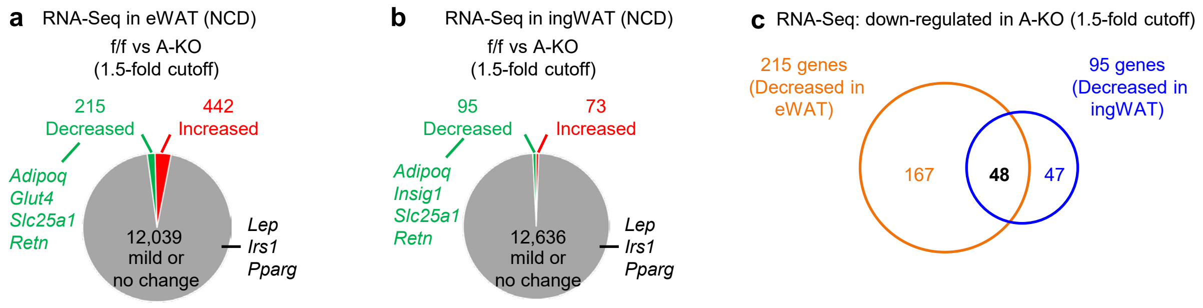


**d** GO analysis: 48 decreased genes in eWAT and iWAT (top terms)

| GO term | Count | P-Value | Gene list |
|---|---|---|---|
| fat cell differentiation | 6 | 1.90E-05 | *Adipoq, Adrb3, Aldh6a1, Fgf10, Pnpla3, Retn* |
| lipid metabolic process | 11 | 2.20E-04 | *Adhfe1, Adipoq, Adora1, Decr1, Enpp2, Ltc4s, Mogat1, Pik3cb, Pnpla3, Proca1, Tmem43* |
| positive regulation of cold-induced thermogenesis | 4 | 1.20E-03 | *Adipoq, Adrb3, Cmklr1, Decr1* |
| regulation of cell migration | 9 | 1.30E-03 | *Acvr1c, Adipoq, Adora1, Cmklr1, Enpp2, Fgf10, Pik3cb, Retn, Srpx2* |
| regulation of cell motility | 9 | 1.90E-03 | *Acvr1c, Adipoq, Adora1, Cmklr1, Enpp2, Fgf10, Pik3cb, Retn, Srpx2* |

**e** RNA-Seq (RPKM): **NCD**

Red: f/f vs A-KO >1.5 fold

| | | eWAT | | | | ingWAT | | | |
|---|---|---|---|---|---|---|---|---|---|
| | | f/f-1 | f/f-2 | A-KO-1 | A-KO-2 | f/f-1 | f/f-2 | A-KO-1 | A-KO-1 |
| Adipogenesis marker | *Pparg* | 121.4 | 99.6 | 95.6 | 85.2 | 106.7 | 71.7 | 87.2 | 64.0 |
| | *Cebpa* | 384.9 | 376.5 | 307.5 | 310.8 | 401.4 | 307.4 | 358.4 | 196.2 |
| | *Cebpb* | 41.7 | 37.7 | 42.1 | 49.0 | 72.2 | 45.4 | 81.0 | 50.8 |
| | *Fabp4* | 13689.3 | 11866.7 | 11922.3 | 13877.3 | 13924.1 | 8873.0 | 7833.5 | 9119.0 |
| Insulin response | *Adipoq* | 2033.7 | 1772.0 | 824.4 | 862.1 | 1214.8 | 1020.2 | 427.1 | 427.3 |
| | *Glut4* | 53.1 | 49.9 | 26.3 | 31.9 | 54.5 | 62.4 | 33.9 | 51.7 |
| | *Insig1* | 39.7 | 43.1 | 46.7 | 26.5 | 65.7 | 38.7 | 23.4 | 17.3 |
| | *Retn* | 1387.5 | 1407.7 | 456.4 | 373.9 | 986.2 | 705.8 | 334.7 | 151.3 |
| Lipogenic TFs | *Lxra* | 29.8 | 29.2 | 28.0 | 30.8 | 35.3 | 29.2 | 49.9 | 26.7 |
| | *Lxrb* | 41.1 | 36.3 | 33.1 | 35.6 | 44.5 | 32.4 | 46.5 | 23.5 |
| | *Srebp1* | 83.8 | 66.2 | 57.7 | 57.2 | 103.4 | 60.2 | 81.3 | 42.4 |
| | *Srebp2* | 12.7 | 14.4 | 16.0 | 14.1 | 15.4 | 14.3 | 21.7 | 12.2 |
| | *Chrebp* | 17.5 | 16.3 | 13.9 | 14.0 | 22.7 | 20.5 | 22.6 | 16.0 |
| Lipogenesis | *Slc25a1* | 164.8 | 135.1 | 75.8 | 61.2 | 171.4 | 139.7 | 67.3 | 47.7 |
| | *Fasn* | 319.7 | 352.8 | 317.7 | 180.4 | 304.3 | 303.1 | 395.3 | 142.1 |
| | *Scd1* | 4024.3 | 3308.2 | 4240.8 | 4318.9 | 2501.8 | 2573.1 | 2518.1 | 2923.7 |
| | *Agpat2* | 171.2 | 188.4 | 160.9 | 94.1 | 216.2 | 163.8 | 136.4 | 51.2 |
| | *Bscl2* | 100.4 | 90.9 | 66.0 | 58.7 | 127.0 | 77.6 | 98.0 | 43.6 |
| Lipid uptake | *Lpl* | 1874.0 | 1928.2 | 2733.7 | 2493.3 | 1142.0 | 1223.7 | 1272.6 | 1291.3 |
| | *Cd36* | 728.2 | 717.5 | 809.3 | 763.2 | 481.0 | 506.1 | 296.5 | 411.4 |
| | *Fatp1* | 189.1 | 99.1 | 118.1 | 133.9 | 251.7 | 119.3 | 163.2 | 109.5 |

**Figure S3. RNA-Seq analysis in WAT of mice with adipocyte-specific deletion of *Hira* under normal chow diet**

All data were from 23-week-old *Hira$^{f/f}$* (f/f) and *Hira$^{f/f}$;Adipoq-Cre* (A-KO) male mice fed with a normal chow diet. Total RNA isolated from two to three mice per genotype was combined in equal amounts into one sample. Two independent samples (biological replicates) for each genotype were used for RNA-seq analysis. **a-b**. RNA-Seq analysis of eWAT (**a**) and ingWAT (**b**). Differentially expressed genes were identified using DESeq2 (8) in R (v3.5.3), applying a threshold of 1.5-fold change and a *P*-value < 0.01. NCD: normal chow diet. **c**, Venn diagrams depicting down-regulated genes in adipose tissues of A-KO mice. **d,** Gene Ontology (GO) analysis of genes defined in **c**. **e**, Expression levels of representative genes are shown in RPKM values of RNA-Seq data.

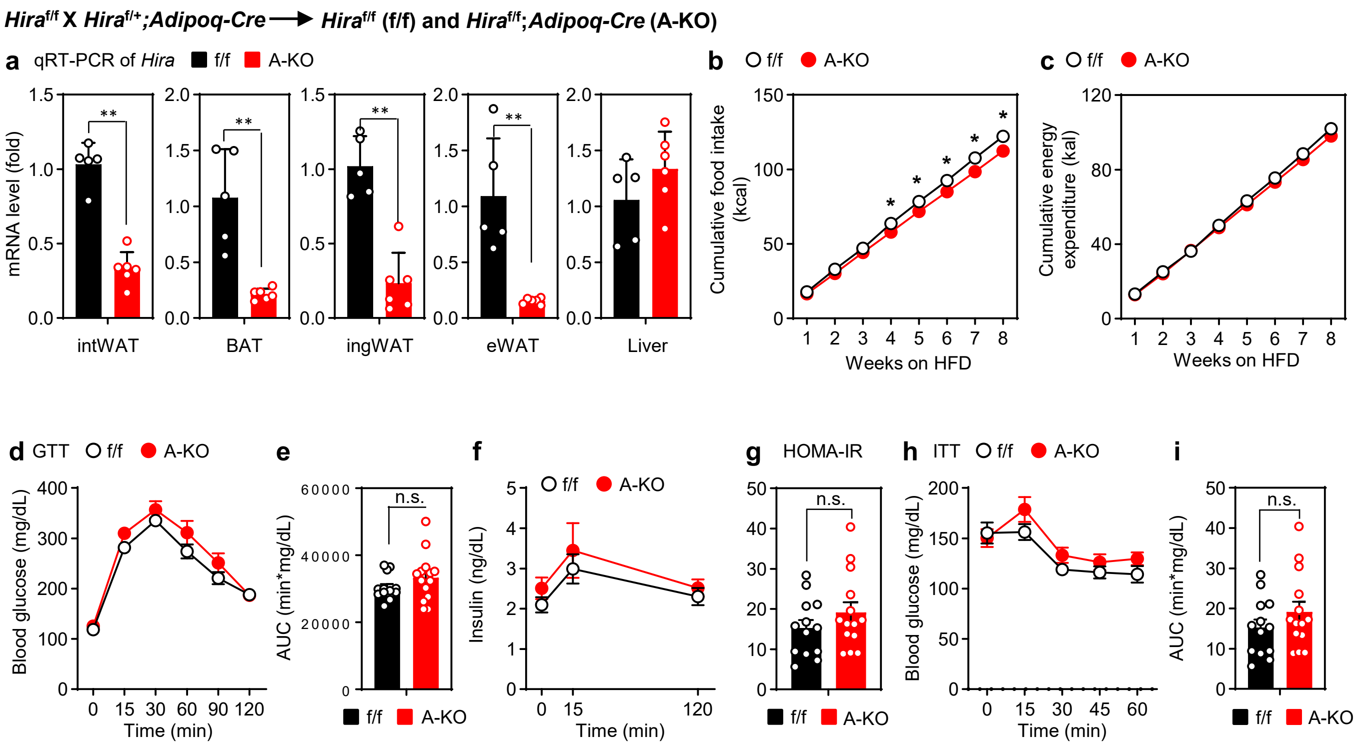


**Figure S4. Characterization of mice with adipocyte-specific deletion of *Hira* during HFD-induced obesity**

Male *Hira*$^{f/f}$ (f/f) and *Hira*$^{f/f}$;AdipoqCre (A-KO) mice (**a-c**, *n* = 5~6 per group. **d-i**, *n* = 13~14 per group) were fed with HFD from the eighth week of age. **a**, qRT-PCR analysis of *Hira* mRNA levels in different tissues. **b-c**, Cumulative food intake (**b**) and cumulative energy expenditure (**c**). **d-f**, Glucose tolerance test (GTT): blood glucose levels (**d**), area under the curve (AUC, **e**) and insulin levels (**f**). **g**, Insulin sensitivity was determined by HOMA-IR. **h-i**, Insulin tolerance test (ITT): blood glucose levels (**h**) and area under the curve (AUC, **i**). All quantitative data for mice are presented as means ± SEM. Statistical comparison between groups was performed using Student's *t*-test (**a**, **b**, **e**, **g** and **i**) or two-way repeated measures ANOVA analysis (**d, f** and **h**). (∗) $P < 0.05$, (∗∗) $P < 0.01$, (∗∗∗) $P < 0.001$.

***Hira*$^{f/f}$ X *Hira*$^{f/+}$*;Adipoq-Cre* ⟶ *Hira*$^{f/f}$ (f/f) and *Hira*$^{f/f}$*; Adipoq-Cre* (A-KO)**

**a** GO analysis of 175 decreased genes (top terms)

| GO Term | Count | Gene list |
|---|---|---|
| organophosphate metabolic process | 25 | *Adcy5, Agpat2, Aldh1l1, Cyb5r4, Dera, Ehhadh, Fasn, G6pdx, Gpd1, Lpgat1, Me1, Mfsd8, Pcx, Pgd, Pik3cb, Pnpla3, Pnpla8, Prps1, Slc25a10, Slc27a1, Slc4a7, Tpi1, Ttc7b, Umps, Vac14* |
| NADP metabolic process | 7 | *Aldh1l1, Cyb5r4, G6pdx, Me1, Pcx, Pgd, Prps1* |
| lipid metabolic process | 28 | *Abhd15, Acad9, Adipoq, Adipor2, Adora1, Agpat2, Bscl2, Decr1, Ehhadh, Fads3, Fasn, G6pdx, Gpd1, Insig1, Lipf, Lpgat1, Ltc4s, Mfsd8, Pcx, Pik3cb, Pnpla3, Pnpla8, Retsat, Slc27a1, Tmem86a, Ttc7b, Vac14, Vldlr* |
| nicotinamide nucleotide metabolic process | 10 | *Aldh1l1, Cyb5r4, G6pdx, Gpd1, Me1, Mfsd8, Pcx, Pgd, Prps1, Tpi1* |
| pyridine nucleotide metabolic process | 10 | *Aldh1l1, Cyb5r4, G6pdx, Gpd1, Me1, Mfsd8, Pcx, Pgd, Prps1, Tpi1* |
| carboxylic acid metabolic process | 22 | *Acad9, Adipoq, Adipor2, Aldh1l1, Ass1, Csad, Decr1, Ehhadh, Fads3, Fasn, Lipf, Ltc4s, Me1, Mfsd8, Npl, Pcx, Pgd, Pnpla3, Pnpla8, Slc27a1, Tph2, Tpi1* |
| oxoacid metabolic process | 22 | *Acad9, Adipoq, Adipor2, Aldh1l1, Ass1, Csad, Decr1, Ehhadh, Fads3, Fasn, Lipf, Ltc4s, Me1, Mfsd8, Npl, Pcx, Pgd, Pnpla3, Pnpla8, Slc27a1, Tph2, Tpi1* |
| phosphate-containing compound metabolic process | 31 | *Acp5, Adcy5, Agpat2, Aldh1l1, Cyb5r4, Dera, Ehhadh, Ephb2, Fam20c, Fasn, G6pdx, Gnptab, Gpd1, Hipk2, Lpgat1, Me1, Mfsd8, Pcx, Pgd, Pik3cb, Pnpla3, Pnpla8, Prps1, Slc25a10, Slc27a1, Slc4a7, Tpi1, Ttc7b, Uhmk1, Umps, Vac14* |

**b** RNA-Seq (RPKM): HFD

Red: f/f vs A-KO >1.5 fold

| | | eWAT | | | | ingWAT | | | |
|---|---|---|---|---|---|---|---|---|---|
| | | f/f-1 | f/f-2 | A-KO-1 | A-KO-2 | f/f-1 | f/f-2 | A-KO-1 | A-KO-2 |
| Adipogenesis marker | *Pparg* | 49.6 | 45.9 | 37.9 | 45.4 | 60.6 | 64.0 | 39.7 | 62.5 |
| | *Cebpa* | 226.8 | 206.8 | 204.3 | 226.2 | 282.4 | 292.9 | 194.6 | 304.9 |
| | *Cebpb* | 50.4 | 44.1 | 56.8 | 45.1 | 52.8 | 47.1 | 47.2 | 65.3 |
| | *Fabp4* | 16322.8 | 10894.2 | 12834.1 | 12535.4 | 21095.5 | 20431.5 | 9878.7 | 17475.7 |
| Insulin response | *Adipoq* | 1302.6 | 1057.9 | 467.7 | 614.5 | 1875.5 | 1840.1 | 387.7 | 603.5 |
| | *Glut4* | 21.1 | 23.3 | 14.9 | 20.7 | 40.2 | 37.1 | 23.1 | 28.5 |
| | *Insig1* | 76.5 | 56.7 | 23.2 | 28.8 | 71.3 | 54.0 | 13.4 | 20.7 |
| | *Retn* | 582.8 | 495.1 | 421.7 | 339.1 | 1113.1 | 1116.6 | 190.0 | 323.9 |
| Lipogenic TFs | *Lxra* | 22.6 | 25.6 | 23.5 | 20.6 | 22.6 | 23.3 | 17.6 | 22.1 |
| | *Lxrb* | 47.1 | 40.8 | 25.5 | 27.7 | 40.0 | 43.8 | 21.3 | 31.9 |
| | *Srebp1* | 34.4 | 39.4 | 38.7 | 40.5 | 40.7 | 45.1 | 23.9 | 34.2 |
| | *Srebp2* | 9.3 | 9.8 | 10.3 | 8.4 | 6.2 | 7.7 | 5.7 | 7.8 |
| | *Chrebp* | 4.6 | 5.3 | 3.7 | 5.3 | 6.7 | 6.6 | 5.4 | 7.0 |
| Lipogenesis | *Slc25a1* | 216.6 | 132.4 | 57.2 | 59.2 | 270.7 | 214.9 | 40.2 | 65.9 |
| | *Fasn* | 184.5 | 168.5 | 60.0 | 87.0 | 140.7 | 127.3 | 45.0 | 71.6 |
| | *Scd1* | 3801.7 | 3677.6 | 3032.0 | 4908.9 | 4255.3 | 4154.1 | 3432.0 | 4977.7 |
| | *Agpat2* | 199.9 | 187.2 | 75.2 | 97.7 | 216.7 | 249.2 | 65.3 | 87.0 |
| | *Bscl2* | 151.7 | 133.4 | 95.4 | 58.5 | 158.1 | 159.4 | 43.5 | 74.9 |
| Lipid uptake | *Lpl* | 1826.5 | 1502.6 | 1259.7 | 1437.9 | 2082.6 | 2069.0 | 1247.0 | 1965.4 |
| | *Cd36* | 731.2 | 672.2 | 287.1 | 407.5 | 782.7 | 810.3 | 321.3 | 426.0 |
| | *Fatp1* | 136.6 | 130.1 | 72.9 | 76.3 | 207.8 | 204.0 | 82.4 | 121.4 |

**Figure S5. Adipocyte-specific deletion of *Hira* reduces expression of *Adipoq* and lipid metabolism genes in WAT during HFD-induced obesity**

All data were from *Hira*$^{f/f}$ (f/f) and *Hira*$^{f/f}$;*Adipoq-Cre* (A-KO) mice ($n$ = 5~6 per group) fed with HFD for 8 weeks from the eighth week of age. **a**, List of all genes associated with GO terms shown in Fig. 3e. **b**, Expression levels of representative genes are shown in RPKM values of RNA-Seq data.

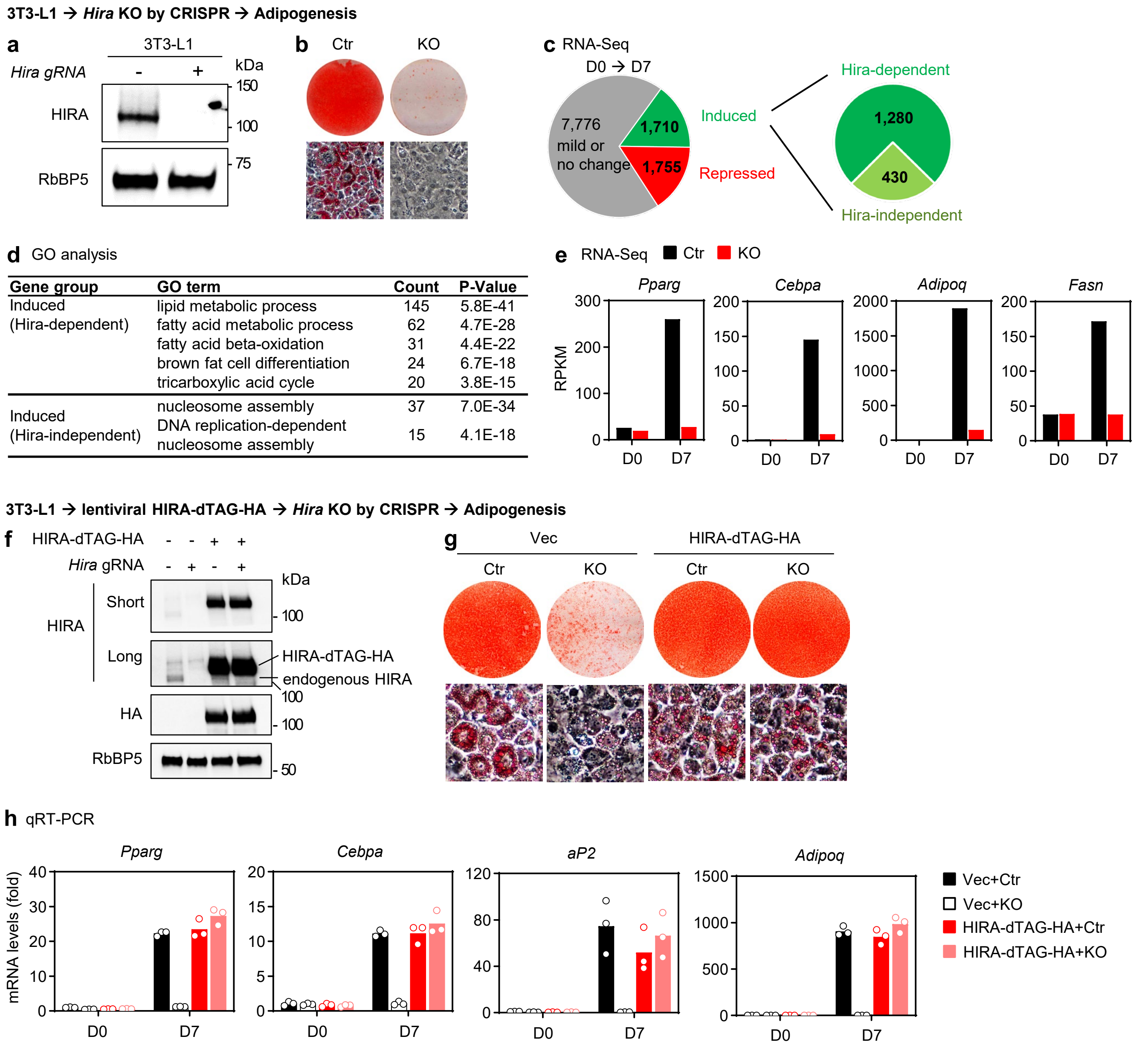


**Figure S6. Knockout of *Hira* in preadipocytes prevents adipogenesis**

**a-e**, 3T3-L1 white preadipocytes were infected with a lentiviral CRISPR/Cas9-*Hira* gRNA to delete *Hira*, followed by adipogenesis assay and RNA-Seq analysis. **a**, Western blot analysis using antibodies indicated on the left. RbBP5 was used as a loading control. **b**, Oil Red O staining at day 7 (D7) of adipogenesis is shown. **c**, RNA-Seq analysis at D7 of adipogenesis. The cutoff for differential expression is 2-fold. **d**, GO analysis of gene groups defined in **c**. **e,** Expression levels of *Pparg*, *Cebpa*, *Adipoq* and *Fasn* at day 0 (D0) and D7 determined by RNA-Seq ($n$ = 1). **f-h**, 3T3-L1 white preadipocytes were infected with lentiviruses expressing HIRA-dTAG-HA, followed by lentiviral CRISPR/Cas9-*Hira* gRNA infection to delete endogenous *Hira*. Adipogenesis assay and qRT-PCR analysis were then performed. **f**, Western blot analyses using antibodies indicated on the left. RbBP5 was used as a loading control. **g**, Oil Red O staining at D7 of adipogenesis. **h**, qRT-PCR analysis of adipogenesis marker genes ($n$ =3).

**3T3-L1 + HIRA-dTAG-HA + *Hira* KO → Adipogenesis → dTAG-13 treatment at D4 or D7 → Western blot & ChIP-Seq**

**a** HIRA binding sites associated genes

| Cluster | Top Go Term | Gene list |
|---|---|---|
| Cluster D | cellular response to insulin stimulus | *Akt1, Akt2, Akt3, Apc, Apobec1, Aprt, Baiap2, Baiap2l2, Bcar1, C2cd5, Cav2, Ccl2, Cdk4, Cpeb2, Dennd4c, Eef2k, Eif4ebp1, Enpp1, Errfi1, Foxc2, Foxo1, Gclc, Ghrhr, Ghsr, Gpld1, Grb2, Gsk3b, Hdac9, Igf1r, Inhbb, Inpp5k, Insr, Irs1, Irs2, Irs3, Irs4, Kat2b, Kl, Lpin1, Mars, Myo5a, Ndel1, Parp1, Pde3b, Pdk4, Pdpk1, Phip, Pik3r1, Pik3r2, Pik3r3, Pkm, Plcb1, Pparg, Ppat, Prkci, Prkdc, Pten, Ptpn1, Ptpn2, Ptpra, Rab10, Rab13, Rab8a, Rhoq, Sgk1, Sh2b2, Shc1, Sik2, Slc25a33, Slc2a4, Slc2a8, Smarcc1, Socs7, Soga1, Sorbs1, Srebf1, Srsf3, Stxbp4, Tbc1d4, Tsc2, Tusc5, Vamp2, Wdr11, Ywhag, Zfp106, Zfp36l1* |
| Cluster E | cellular response to insulin stimulus | *Adipoq, Akt1, Akt3, Apc, Apobec1, Baiap2, Baiap2l1, Bcar1, C2cd5, Cav2, Cpeb2, Dennd4c, Dnaic1, Eef2k, Eif4ebp2, Enpp1, Errfi1, Fer, Foxc2, Foxo1, Gclc, Ghsr, Grb2, Gsk3a, Gsk3b, Hdac9, Igf1r, Igfbp1, Inhbb, Insr, Irs2, Irs3, Irs4, Kat2b, Kl, Lpin1, Mup4, Myo5a, Ndel1, Parp1, Pde3b, Pdk2, Pdk4, Phip, Pik3r1, Pik3r2, Pik3r3, Pklr, Plcb1, Pparg, Ppat, Prkci, Pten, Ptpn1, Ptpn2, Rab10, Rab8a, Rhoq, Rpe65, Rps6kb1, Sgk1, Sh2b2, Sik2, Slc25a33, Slc2a4, Slc2a8, Slc9a1, Sorbs1, Srebf1, Srsf3, Tbc1d4, Trib3, Tusc5, Uchl3, Ucp2, Usf1, Vamp2, Wdr11, Wdtc1, Ywhag, Zfp36l1* |
| | fatty acid metabolic process | *Aacs, Abcd2, Abcd3, Abhd5, Acaa2, Acaca, Acacb, Acadl, Acadm, Acads, Acadvl, Acat2, Acat3, Acod1, Acot1, Acot12, Acot2, Acot3, Acot4, Acot5, Acot6, Acox2, Acox3, Acsbg1, Acsf3, Acsl1, Acsl3, Acsl4, Acsl5, Acsm1, Acsm3, Acsm4, Acss2, Adh7, Adipoq, Adipor2, Alkbh7, Alox15, Alox5ap, Angptl3, Ankrd23, Apoa2, Auh, Bdh2, C3, Cbr4, Cpt1a, Cpt1c, Cpt2, Crat, Crem, Cyb5a, Cyp1b1, Cyp2b10, Cyp2b13, Cyp2c39, Cyp2c44, Cyp2c55, Cyp2c67, Cyp2d22, Cyp2d26, Cyp2f2, Cyp4a12a, Cyp4a32, Cyp4v3, Decr1, Dld, Echdc1, Echdc2, Echs1, Eci1, Eci2, Eci3, Edn1, Edn2, Elovl6, Etfa, Fa2h, Faah, Fabp3, Fads1, Fads2, Fads6, Fasn, Ggt5, Ghr, Gpam, Hacd2, Hacd3, Hacl1, Hpgd, Hsd17b12, Ivd, Lep, Lias, Lipe, Lpin1, Lpin2, Lpl, Lta4h, Ltc4s, Lypla2, Mapk14, Mcat, Mgll, Mgst2, Mgst3, Myo5a, Ndufab1, Paf1, Pccb, Pecr, Per2, Pex13, Pex2, Pex5, Pla2g15, Pnpla3, Pnpla8, Por, Ppara, Ppard, Ppargc1a, Prg3, Prkaa1, Prkaa2, Prkab1, Prkab2, Prkag2, Prkag3, Prkar2b, Ptges, Ptges2, Ptges3, Ptgis, Ptgr1, Ptgr2, Ptgs1, Qk, Scd1, Scd2, Scd3, Scd4, Sesn2, Sgpl1, Slc25a17, Slc27a3, Slc27a4, Snca, Stat5a, Stat5b, Syk, Tbxas1, Tecr, Tecrl, Thnsl2, Tnxb, Ucp3, Zadh2,* |
| Cluster F | cellular response to insulin stimulus | *Adipoq, Akt1, Akt3, Apc, Bcar1, Cav2, Ccl2, Cdk4, Cpeb2, Eef2k, Enpp1, Errfi1, Fer, Foxc2, Foxo1, Ghsr, Gsk3b, Hdac9, Igf1r, Inhbb, Inpp5k, Insr, Irs1, Irs2, Irs3, Kat2b, Kl, Lpin1, Mup16, Mup18, Mup2, Myo5a, Parp1, Pde3b, Pdk4, Pdpk1, Phip, Pik3r1, Pik3r3, Pkm, Plcb1, Pparg, Pten, Ptpn1, Rab10, Rab31, Rab8a, Rpe65, Rps6kb1, Sgk1, Sh2b2, Shc1, Sik2, Slc25a33, Slc2a4, Soga1, Sorbs1, Srebf1, Srsf3, Star, Stxbp4, Tbc1d4, Tusc5, Usf1, Vamp2, Wdr11, Zfp36l1* |

**b** Motif analysis: HIRA$^+$ AEs (17,862) at D4

| Motif | P-Value |
|---|---|
| AP1 | 1E-604 |
| C/EBP | 1E-418 |
| NF1 | 1E-403 |
| TEAD | 1E-188 |
| C/EBP:AP1 | 1E-144 |
| PPAR | 1E-72 |

**c** Motif analysis: HIRA$^+$ AEs (599) associated with insulin response and lipid metabolism genes

| Motif | P-Value |
|---|---|
| C/EBP | 1E-23 |
| AP1 | 1E-21 |
| NF1 | 1E-14 |
| TEAD | 1E-09 |
| EBF | 1E-04 |
| GR | 1E-04 |

**d** Genomic tracks (GSE160605)

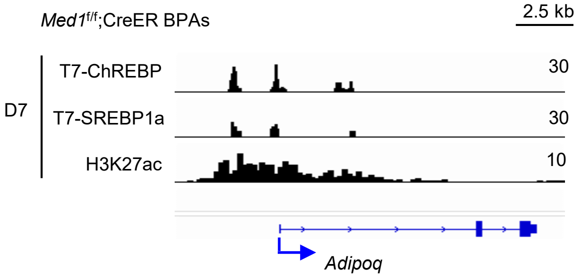


### Figure S7. HIRA directly binds to insulin response and lipid metabolism genes in adipocytes

**a,** List of all genes associated with top GO terms in clusters D, E and F defined in Fig. 4d. **b**, Motif analysis of HIRA$^+$ active enhancers (AEs) at D4. **c**, Motif analysis of HIRA$^+$ AEs associated with insulin response and lipid metabolism genes defined in Fig. 4d. **d**, ChIP-Seq profiles of T7-ChREBP, T7-SREBP1a and H3K27ac on the *Adipoq* gene locus in adipocytes at D7. Data are from GSE160605 (5).

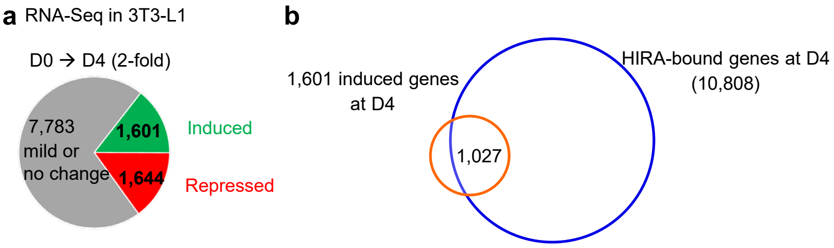


**c** GO analysis (1,027 genes)

| GO term | Count | P-Value |
|---|---|---|
| organic acid metabolic process | 132 | 1.1E-33 |
| oxoacid metabolic process | 130 | 2.7E-33 |
| carboxylic acid metabolic process | 128 | 3.5E-33 |
| lipid metabolic process | 149 | 1.1E-25 |
| monocarboxylic acid metabolic process | 93 | 3.3E-25 |
| small molecule catabolic process | 68 | 2.7E-23 |

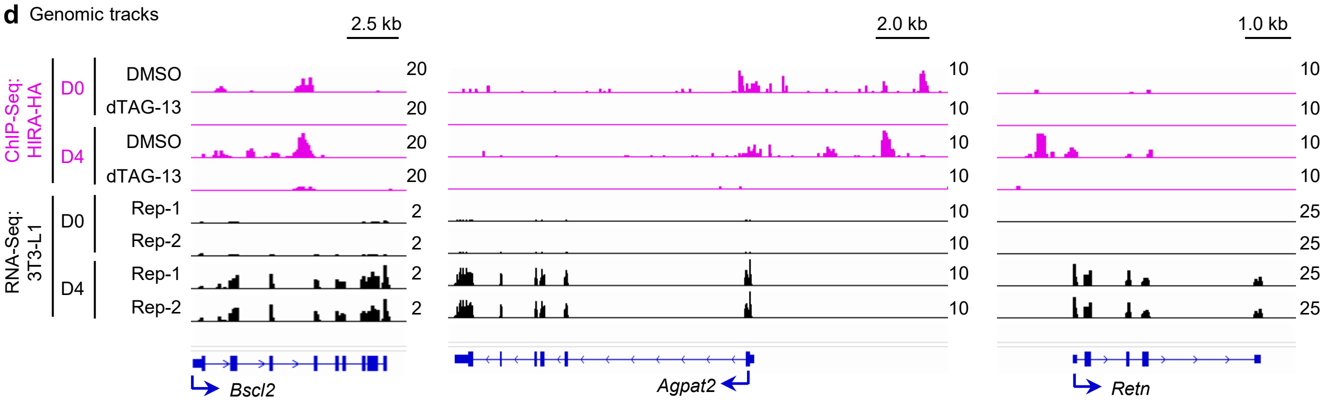


**Figure S8. Identification of genes induced at D4 of adipogenesis and bound by HIRA in 3T3-L1 adipocytes**

**a**, RNA-Seq analysis at D0 and D4 of adipogenesis in 3T3-L1 cells ($n$ = 2). Differentially expressed genes were identified using DESeq2 (8) in R (v3.5.3), applying a threshold of 2-fold change and a $P$-value < 0.01. **b**, Venn diagrams depicting genes that were induced over 2-fold at D4 and bound by HIRA. **c**, GO analysis of genes defined in **b**. **d**, Profiles of HIRA binding and RNA-Seq signals around *Bscl2*, *Agpat2* and *Retn* gene loci.

**a** GO analysis (6 h depletion of HIRA: Q2)

| GO Term | Count | P-Value |
|---|---|---|
| organic acid metabolic process | 41 | 1.10E-13 |
| oxoacid metabolic process | 40 | 3.00E-13 |
| carboxylic acid metabolic process | 39 | 6.50E-13 |
| generation of precursor metabolites and energy | 26 | 4.70E-12 |
| monocarboxylic acid metabolic process | 31 | 1.20E-11 |
| organophosphate metabolic process | 39 | 5.80E-11 |

**b** GO analysis (6 h depletion of HIRA: Q3)

| GO Term | Count | P-Value |
|---|---|---|
| phosphate-containing compound metabolic process | 47 | 5.10E-09 |
| phosphorus metabolic process | 47 | 5.40E-09 |
| generation of precursor metabolites and energy | 21 | 2.70E-08 |
| carboxylic acid metabolic process | 31 | 4.20E-08 |
| oxoacid metabolic process | 31 | 7.20E-08 |
| organic acid metabolic process | 39 | 5.80E-11 |

**c** GO analysis (6 h depletion of HIRA: Q4)

| GO Term | Count | P-Value |
|---|---|---|
| cellular response to oxygen levels | 16 | 1.10E-08 |
| negative regulation of metabolic process | 72 | 1.50E-08 |
| cellular response to hypoxia | 14 | 2.00E-08 |
| cellular response to decreased oxygen levels | 15 | 2.40E-08 |
| positive regulation of metabolic process | 80 | 3.30E-08 |
| regulation of cold-induced thermogenesis | 13 | 1.80E-07 |

**d** GO analysis (24 h depletion of HIRA: Q2)

| GO Term | Count | P-Value |
|---|---|---|
| carboxylic acid metabolic process | 40 | 1.40E-13 |
| oxoacid metabolic process | 40 | 3.00E-13 |
| organic acid metabolic process | 40 | 5.20E-13 |
| generation of precursor metabolites and energy | 26 | 4.70E-12 |
| small molecule catabolic process | 24 | 3.00E-11 |
| phosphate-containing compound metabolic process | 51 | 7.10E-11 |

**e** GO analysis (24 h depletion of HIRA: Q3)

| GO Term | Count | P-Value |
|---|---|---|
| oxoacid metabolic process | 32 | 1.80E-08 |
| organic acid metabolic process | 32 | 2.80E-08 |
| carboxylic acid metabolic process | 31 | 3.80E-08 |
| lipid metabolic process | 36 | 1.70E-06 |
| fatty acid metabolic process | 18 | 4.00E-06 |
| monocarboxylic acid metabolic process | 22 | 4.80E-06 |

**f** GO analysis (24 h depletion of HIRA: Q4)

| GO Term | Count | P-Value |
|---|---|---|
| regulation of apoptotic process | 50 | 2.60E-09 |
| regulation of programmed cell death | 50 | 6.60E-09 |
| negative regulation of metabolic process | 71 | 2.00E-08 |
| positive regulation of metabolic process | 79 | 4.00E-08 |
| negative regulation of biosynthetic process | 60 | 5.00E-08 |
| positive regulation of macromolecule metabolic process | 72 | 2.40E-07 |

**g** GO analysis of genes in Q1 (6 h depletion of HIRA)

| GO Term | Count | Gene list |
|---|---|---|
| lipid metabolic process | 47 | *Abcd2, Abhd11, Acadm, Acat2, Acp6, Acadvl, Adh1, Adhfe1, Adipoq, Aldh3b2, Akr7a5, Amacr, Apoc1, C3, Cav1, Cpt2, Cyp2f2, Cyp39a1, Dgat1, Dgat2, Echdc3, Etfb, Fgfr1, G6pdx, Gk5, Gpd1, Gnpat, Gsta3, Hacd2, Hsd17b10, Lipe, Ltc4s, Mgst2, Mgst3, Mlxipl (Chrebp), Plaat3, Plin1, Pxmp4, Rdh12, Rdh5, Retsat, Scp2, Slc22a4, St3gal1, Sult1a1, Thrsp* |
| organic acid metabolic process | 35 | *Abcd2, Acadm, Acat2, Aco2, Acadvl, Adh1, Adhfe1, Adipoq, Aldh4a1, Amacr, C3, Cs, Cpt2, Cyp2f2, Cyp39a1, Dlst, Echdc3, Etfb, Fgfr1, Gamt, Hacd2, Hpdl, Hsd17b10, Kyat1, Lipe, Ltc4s, Mgst2, Mgst3, Mpst, Mpc2, Pgk1, Scp2, Slc5a6, Stat5a, St3gal1* |
| oxoacid metabolic process | 34 | *Abcd2, Acadm, Acat2, Aco2, Acadvl, Adh1, Adhfe1, Adipoq, Aldh4a1, Amacr, C3, Cs, Cpt2, Cyp2f2, Cyp39a1, Dlst, Echdc3, Etfb, Fgfr1, Gamt, Hacd2, Hpdl, Hsd17b10, Kyat1, Lipe, Ltc4s, Mgst2, Mgst3, Mpst, Mpc2, Pgk1, Scp2, Slc5a6, St3gal1* |
| carboxylic acid metabolic process | 33 | *Abcd2, Acadm, Acat2, Aco2, Acadvl, Adh1, Adhfe1, Adipoq, Aldh4a1, Amacr, C3, Cs, Cpt2, Cyp2f2, Cyp39a1, Dlst, Echdc3, Etfb, Gamt, Hacd2, Hpdl, Hsd17b10, Kyat1, Lipe, Ltc4s, Mgst2, Mgst3, Mpst, Mpc2, Pgk1, Scp2, Slc5a6, St3gal1* |
| regulation of lipid metabolic process | 20 | *Abcd2, Acadvl, Adipoq, Agt, Apoc1, Bscl2, Cav1, Chp1, C3, Cidec, Dgat1, Dgat2, Mlxipl (Chrebp), Pgk1, Phb2, Pdk2, Prkaca, Scp2, Stat5a, Thrsp* |
| regulation of small molecule metabolic process | 20 | *Abcd2, Acadm, Acadvl, Adipoq, Agt, Apoc1, Atp2b4, Cav1, Cd320, Dgat1, Dgat2, Gpd1, Igfbp4, Insr, Mlxipl (Chrebp), Pgk1, Phkg1, Pdk2, Prkaca, Scp2* |

**h** GO analysis of genes in Q1 (24 h depletion of HIRA)

| GO Term | Count | Gene list |
|---|---|---|
| lipid metabolic process | 50 | *Abhd15, Acacb, Acat2, Acadm, Acsl1, Adipor2, Adipoq, Adhfe1, Agpat2, Aldh3b2, Apoc1, C3, Cat, Cav1, Crat, Cyp2f2, Dagla, Dgat1, Dgat2, Dgkd, Dolpp1, Etfb, Gpam, Gpd1, Gnpat, Hacd2, Hsd17b14, Itpka, Lipe, Lpin1, Ltc4s, Mgst3, Mlxipl (Chrebp), Naaa, Pcx, Pla2g12a, Plaat3, Plin1, Ppard, Ptges, Ptges2, Pxmp4, Rdh12, Retsat, Scarb1, Scp2, Slc22a4, Srebf1, Thrsp* |
| organic acid metabolic process | 39 | *Acat2, Acacb, Acadm, Aco1, Aco2, Acsl1, Adhfe1, Adipoq, Adipor2, Adss1, Aldh4a1, C3, Crat, Cyp2f2, Dagla, Etfb, Fmo1, Gamt, Gpam, Grhpr, Hacd2, Hagh, Lipe, Lpin1, Ltc4s, Mgst3, Mpc2, Mtarc2, Mthfd2, Naaa, Nat8l, Pcx, Ppard, Prodh, Ptges, Ptges2, Scp2, Slc5a6, Stat5a* |
| oxoacid metabolic process | 37 | *Acat2, Acacb, Acadm, Aco1, Aco2, Acsl1, Adhfe1, Adipoq, Adipor2, Adss1, Aldh4a1, C3, Crat, Cyp2f2, Dagla, Etfb, Gamt, Gpam, Grhpr, Hacd2, Hagh, Lipe, Lpin1, Ltc4s, Mgst3, Mpc2, Mtarc2, Mthfd2, Naaa, Nat8l, Pcx, Ppard, Prodh, Ptges, Ptges2, Scp2, Slc5a6* |
| carboxylic acid metabolic process | 36 | *Acat2, Acacb, Acadm, Aco1, Aco2, Acsl1, Adhfe1, Adipoq, Adipor2, Adss1, Aldh4a1, C3, Crat, Cyp2f2, Dagla, Etfb, Gamt, Gpam, Grhpr, Hacd2, Hagh, Lipe, Lpin1, Ltc4s, Mgst3, Mpc2, Mthfd2, Naaa, Nat8l, Pcx, Ppard, Prodh, Ptges, Ptges2, Scp2, Slc5a6* |
| monocarboxylic acid metabolic process | 30 | *Acat2, Acacb, Acadm, Acsl1, Adipoq, Adipor2, C3, Crat, Cyp2f2, Dagla, Etfb, Gamt, Gpam, Grhpr, Hacd2, Hagh, Lipe, Lpin1, Ltc4s, Mgst3, Mpc2, Mthfd2, Naaa, Nat8l, Pcx, Ppard, Ptges, Ptges2, Scp2, Slc5a6* |
| regulation of lipid metabolic process | 25 | *Acacb, Adipoq, Agt, Apoc1, Bscl2, C3, Cav1, Chp1, Cidec, Dgat1, Dgat2, Fmo1, Gpam, Mlxipl (Chrebp), Pcx, Pdk2, Ppard, Scarb1, Scp2, Sik2, Slc45a3, Srebf1, Stat5a, Thrsp, Rorc* |

**Figure S9. Data associated with Fig. 5**

**a-c**, GO analysis of Q2 (**a**), Q3 (**b**) and Q4 (**c**) genes following 6 h depletion of HIRA defined in Fig. 5b left. **d-f**, GO analysis of Q2 (**d**), Q3 (**e**) and Q4 (**f**) genes following 24 h depletion of HIRA defined in Fig. 5b right. **g-h**, List of all genes associated with GO terms in Fig. 5c left (**g**) and right (**h**).

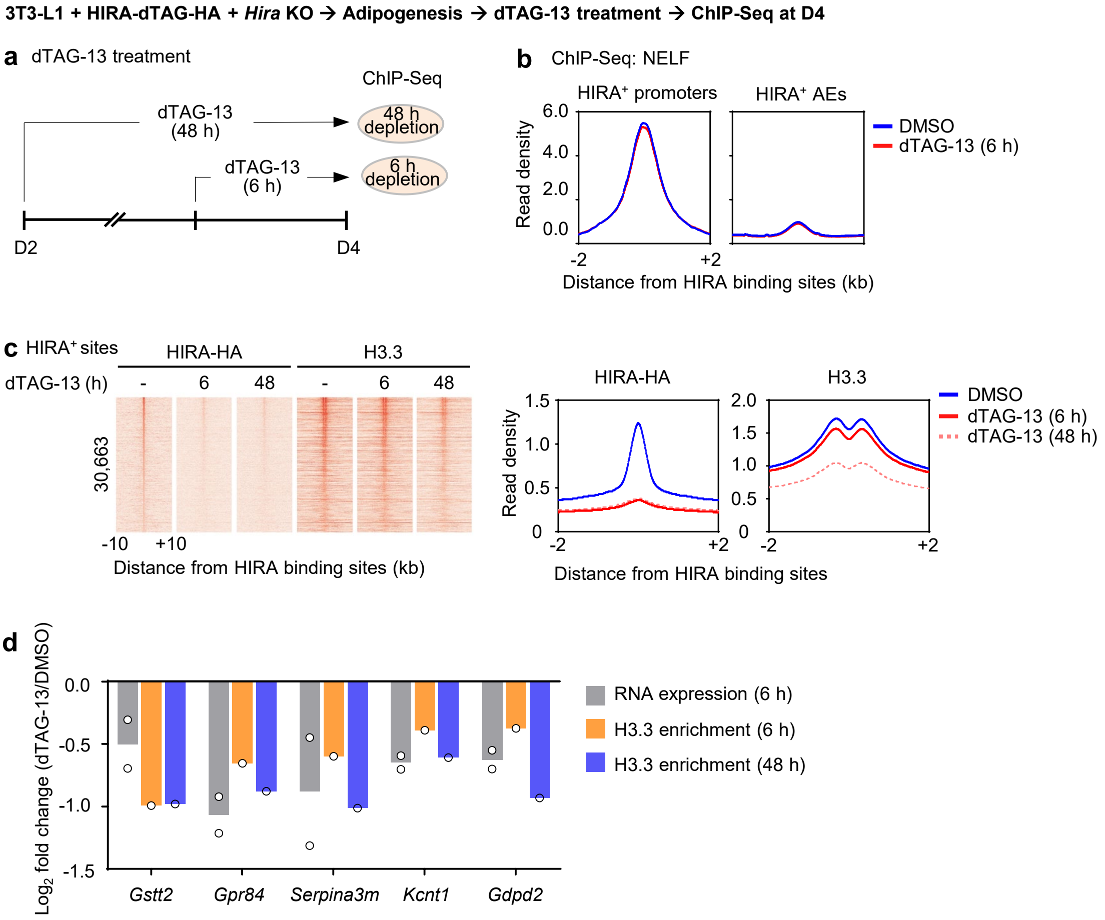


**Figure S10. Acute depletion of HIRA impairs RNA Pol II pause release on insulin response and lipid metabolism genes in adipocytes**

3T3-L1 white preadipocytes were infected with a lentiviral vector expressing HIRA with C-terminal dTAG and HA double tags, followed by lentiviral CRISPR/Cas9-*Hira* gRNA to delete endogenous *Hira*. dTAG-13 treatment was done at D4, followed by ChIP-Seq analysis. **a**, Schematic of ChIP-Seq analysis following HIRA depletion for 6 h or 48 h. **b**, Average binding profiles of NELF around the center of HIRA$^+$ promoters and AEs after 6 h depletion of HIRA. **c,** Heat maps (left) and average profiles (right) showing HIRA-HA binding and H3.3 enrichment aligned to the center of all HIRA$^+$ sites. **d**, Bar graph showing representative gene transcription changes upon 6 h depletion of HIRA (gray) and H3.3 enrichment on TSS $\pm$ 2 kb upon 6 h (orange) or 48 h (blue) depletion of HIRA.

### Materials and Methods

#### Plasmids, antibodies, and chemicals

Lentiviral plasmid pLV-EF1a-IRES-hygro–HIRA-dTAG-HA was generated by following steps: (1) dTAG-HA (FKBP_F36V-HA) fragment (Addgene #91796) was C-terminally subcloned into lentiviral vector pLV-EF1a-IRES-hygro (Addgene #85134) to generate pLV-EF1a-IRES-hygro-dTAG-HA. (2) Human *HIRA* was PCR-amplified using cDNA from 293T cells as templates and was subcloned into pLV-EF1a-IRES-hygro-dTAG-HA. (3) Q5 Site-Directed Mutagenesis Kit (NEB #E0554S) was used to mutate gRNA sequence in the plasmid. Plasmid was confirmed by DNA sequencing. Anti-HIRA (04-1488) and anti-MED1 (17-10530) antibodies were from Sigma. Anti-HA (3724S) for western blot was from Cell Signaling Technology. Anti-HA (13-2010), anti-H3K4me1 (13-0040) and anti-H3.3 (13-0061) for ChIP-Seq were from EpiCypher. Anti-H3K27ac (ab4729) was from Abcam. Anti-RbBP5 (A300-109A) and anti-BRD4 (A301-985A50) were from Bethyl Laboratories. Anti-Pol II-NTD (14958S), anti-S5P-Pol II (13523), anti-S2P-Pol II (13499S), anti-CDK9 (2316T), anti-SPT6 (15616), anti-NELF (12265S) and anti-CBP (7389S) were from Cell Signaling Technology. 500 nM dTAG-13 (Biotechne #6605) was used for depletion of HIRA-dTAG-HA.

#### Generation of mouse strains

*Hira*$^{f/f}$ mice (1) were crossed with *Adipoq-Cre* (Jackson 028020) to generate *Hira*$^{f/f}$;*Adipoq-Cre*. For genotyping *Hira* alleles, PCR was performed using the following primers: 5′-TGCTAAAGAAAACTAGCCGAG-3′ and 5′-TGTGTTTGGTACCCCACAAC-3′. PCR amplified 190 bp from the wild-type and 300 bp from the floxed allele.

All mouse experiments were performed in accordance with the NIH Guide for the Care and Use of Laboratory Animals and approved by the Animal Care and Use Committee of the National Institute of Diabetes and Digestive and Kidney Diseases, NIH.

#### Metabolic studies

For determination of serum metabolites, blood was collected from the tail vein of mice fed with standard laboratory mouse chow (13.6% calories from fat, 60% calories from carbohydrate, 26.4% calories from protein; LabDiet 5018) or high-fat diet (60% calories from fat, 20% calories from carbohydrate, 20% calories from protein; Research Diets D12492). Serum was obtained by centrifuging blood samples at 12,000*g* for 4 min at room temperature. Serum insulin and leptin concentrations were measured by an ELISA kit from CrystalChem and R&D Systems, respectively. Serum concentration of free fatty acid (Fujifilm), free glycerol (Sigma), total triglyceride (Pointe Scientific Inc.), and total cholesterol (Pointe Scientific Inc.) were measured using the indicated calorimetric assay. For glucose tolerance tests (GTT), mice were fasted overnight for 16 h. For insulin tolerance tests (ITT), mice were fasted for 4 h. Glucose (1 g/kg) or human insulin (0.75 U/kg Humulin, Eli Lilly), respectively, were administered intraperitoneally (i.p.), and blood was collected from the tail vein at specific time points. For both tests, blood glucose levels were determined using a portable glucometer (Contour Glucometer, Bayer). HOMA-IR (Homeostatic model assessment for insulin resistance) index was calculated as previously described (2). CL316,243 (0.1 mg/kg; Sigma-Aldrich) or saline were administered i.p. and $O_2$ consumption was measured at 30°C from 1 to 4 h after injection. For in vivo lipolysis, blood was collected 20 min after the injection of CL316,243 (0.1 mg/kg i.p.) or saline.

Body composition was measured with the EchoMRI 3-in-1 analyzer (Echo Medical Systems). In singly housed mice fed with HFD, body composition and food intake were measured weekly over the course of 8 weeks, and weekly energy expenditure was calculated by energy balance technique (3).

**HIRA protein depletion in 3T3-L1 white preadipocytes and adipogenesis assay**

To knock out *Hira* in 3T3-L1 white preadipocytes, cells were infected with lentiviral CRISPR/Cas9 gRNA (TGTGTGCGGTGGTCAAACAG) targeting *Hira*, followed by selection with 2 μg/mL puromycin for 6 days.

To enable temporal control of HIRA protein depletion, 3T3-L1 preadipocytes were infected with lentiviruses pLV-EF1a-IRES-hygro–HIRA-dTAG-HA. After selection with 200 μg/mL hygromycin for 12 days, cells were infected with lentiviral CRISPR/Cas9 *Hira* gRNA.

For the adipogenesis assay, cells were plated in growth medium (DMEM plus 10% bovine serum 4 days before the induction and were induced with 10 μg/mL insulin, 0.5 mM IBMX and 1 μM DEX in fetal bovine serum (FBS) medium (DMEM plus 10% FBS) for 2 days. After 2 days of induction, the culture medium was changed to DMEM supplemented with FBS and insulin only (4).

**Western blot and qRT-PCR**

Western blot of nuclear extracts was done as described (5). Total RNA was extracted using TRIzol (Invitrogen) and reverse-transcribed using a ProtoScript II first strand cDNA synthesis kit (NEB E6560L), following the manufacturer's protocols. qRT-PCR SYBR primers are shown in Supplementary Table 2.

**RNA-Seq library preparation**

Total RNA (1 μg) was subjected to the NEBNext poly(A) mRNA magnetic isolation module (NEB E7490L) to isolate mRNA and proceeded directly to double-stranded cDNA synthesis. Library construction was done using the NEBNext Ultra II RNA library preparation kit for Illumina (NEB E7770L) following the manufacturer's protocol.

**Nascent RNA-Seq library preparation**

Differentiating cells (D4) were labeled with 0.5 mM ethylene uridine (EU) for 30 min before collection. After RNA extraction, 1 μg of total RNA was subjected to the depletion of rRNA using the NEBNext rRNA depletion kit (NEB E7405L). To capture nascent RNA, the sample was biotinylated by the Click-iT nascent RNA capture kit (ThermoFisher C10365) according to the manufacturer's instructions. Captured RNA was reverse transcribed into cDNA using SuperScript™ VILO™ cDNA Synthesis Kit (Thermo, 11754050) and subjected to library construction using the NEBNext Ultra™ II RNA Library Prep Kit for Illumina (NEB, E7770) following the manufacturer's protocol. Sequencing libraries were analyzed with Qubit and pooled and sequenced on NovaSeq X.

**ChIP-Seq library preparation**

ChIP-Seq was performed as described in detail previously (6). ChIP-Seq was done in the presence of *Drosophila* spike-in chromatin and antibody following the manufacturer's protocol (Active Motif). ChIP-Seq library construction was done using a NEBNext Ultra II DNA library preparation kit for Illumina (NEB E7645L) following the manufacturer's protocol. Sequencing libraries were analyzed with Qubit and pooled and sequenced on a NovaSeq 6000, NovaSeq X, or NextSeq 2000.

**Computational analysis**

### Steady-state and nascent RNA-Seq data analysis

Sequencing data were aligned to the mouse mm9 reference genome using the STAR software (7) with default parameters. For steady-state RNA-Seq, aligned reads on exons were calculated to determine the reads per kilobase per million (RPKM) as a measurement of gene expression. Genes with RPKM values exceeding 5 were considered as expressed. For nascent RNA-Seq, read counts were calculated over gene bodies (from transcription start site to transcription end site) to reflect active transcription. Differentially expressed genes were identified using DESeq2 (8) in R (v3.5.3), applying a threshold of 1.5-fold change and a $P$-value < 0.01. To identify the functional significance of differentially expressed genes, Gene ontology (GO) analysis was performed using the DAVID tool (9) (https://davidbioinformatics.nih.gov/home.jsp).

### ChIP-Seq peak calling

Sequencing data were aligned to the mouse mm9 reference genome using Bowtie2. For the identification of ChIP-Seq enriched regions, SICER (version 2) (10) was used. For HIRA-HA and SPT6, a window size of 50 bp, gap size of 50 bp, and a false discovery rate of 1E-10 were used. For S5P-Pol II, S2P-Pol II, and CDK9, a window size of 50 bp, gap size of 50 bp, and a false discovery rate of 1E-3 were used. For H3K27ac, a window size of 200 bp, gap size of 200 bp, and a false discovery rate of 1E-3 were used. The effective genome fraction of 0.8 was applied to all ChIP-Seq data.

### Normalization of ChIP-Seq data

To assess the Spike-in chromatin incorporated into the ChIP reaction, sequencing data were aligned to the drosophila dm6 reference genome using Bowtie2. For data normalization, the sample with the fewest drosophila reads was first identified. Then, the normalization factor was determined by dividing the drosophila reads of the sample with the lowest count by those of other samples. After calculation, each sample's reads were scaled by multiplying them with their respective normalization factor to generate heatmap matrices or profiles.

### Pausing index calculation and cumulative distribution analysis

Pausing index values were calculated for each gene as the ratio of Pol II read density in the promoter-proximal region (−100 bp to +300 bp relative to the TSS) to the read density in the gene body (+300 bp from the TSS to +3 kb beyond the transcription end site [TES]). For each condition (DMSO and dTAG-13), the resulting pausing index values were used to assess overall distribution trends. To generate a normalized cumulative distribution, we calculated the cumulative probability of each pausing index value based on the mean and standard deviation of the dataset. These cumulative probabilities were then used to visualize the relative distribution of pausing indices under each condition.

### Data availability

RNA-Seq and ChIP-Seq data sets generated in the paper have been deposited in NCBI Gene Expression Omnibus under accession number GSE288013.

**Supplemental Table 1: 48 decreased genes in eWAT and ingWAT under normal chow diet**

| Entrez ID | Gene name | eWAT | | | | ingWAT | | | |
|---|---|---|---|---|---|---|---|---|---|
| | | f/f-1 | f/f-2 | A-KO-1 | A-KO-2 | f/f-1 | f/f-2 | A-KO-1 | A-KO-2 |
| 269275 | *Acvr1c* | 22.6 | 22.3 | 15.3 | 13.9 | 12.7 | 12.4 | 5.1 | 7.3 |
| 76187 | *Adhfe1* | 41.6 | 46.1 | 23.7 | 21 | 25.3 | 36.3 | 13.1 | 18.3 |
| 11450 | *Adipoq* | 2033.7 | 1772 | 824.4 | 862.1 | 1214.8 | 1020.2 | 427.1 | 427.3 |
| 11539 | *Adora1* | 15.7 | 13.5 | 5.5 | 5 | 20 | 18.5 | 9.5 | 6.6 |
| 11556 | *Adrb3* | 291 | 222 | 87.5 | 107.2 | 193.7 | 135.9 | 44.9 | 34.2 |
| 231510 | *Agpat9* | 39.8 | 34.8 | 21.8 | 20.4 | 31.7 | 22.3 | 11 | 10.6 |
| 104776 | *Aldh6a1* | 188.5 | 159.2 | 96.5 | 100.2 | 128.4 | 125.4 | 59.2 | 84.7 |
| 109979 | *Art3* | 168.1 | 131 | 66.5 | 83 | 121.4 | 122.1 | 62.3 | 71.6 |
| 12038 | *Bche* | 32.7 | 28.1 | 9.3 | 10.3 | 12.3 | 15.3 | 3.2 | 6.9 |
| 11537 | *Cfd* | 8377.3 | 5332 | 3204.6 | 3516.4 | 10220.8 | 5108.4 | 4326 | 2282.1 |
| 14747 | *Cmklr1* | 72.1 | 62.2 | 36.9 | 49.3 | 49.9 | 54.5 | 30.6 | 26.9 |
| 67460 | *Decr1* | 52.3 | 53.4 | 28 | 27.6 | 69.4 | 51.4 | 34.8 | 28.6 |
| 18606 | *Enpp2* | 114.4 | 96.8 | 56.9 | 66.9 | 76.6 | 68.1 | 28.5 | 36.5 |
| 14165 | *Fgf10* | 11.3 | 9.2 | 4.8 | 3.9 | 13.6 | 9.1 | 4.5 | 3.9 |
| 14265 | *Fmr1* | 32.3 | 56.5 | 15.1 | 16.3 | 17.8 | 31.6 | 6.6 | 12.2 |
| 244757 | *Glb1l2* | 104.2 | 114.7 | 48.5 | 51.2 | 48.2 | 55.3 | 20.4 | 31.6 |
| 15426 | *Hoxc8* | 51.5 | 50.8 | 34.7 | 29.4 | 27.1 | 32.2 | 13.6 | 16.8 |
| 76933 | *Ifi27l2a* | 1155.4 | 652.7 | 131.4 | 107 | 1936.7 | 1246.4 | 223.9 | 95.7 |
| 66307 | *Isoc1* | 50.5 | 54.1 | 27.8 | 29.9 | 33.3 | 35.3 | 10.2 | 16.9 |
| 238076 | *Kcns3* | 12.8 | 16.3 | 5.1 | 9.2 | 7.5 | 9.3 | 3.7 | 4.9 |
| 77113 | *Klhl2* | 69.1 | 50.7 | 31.2 | 34.2 | 45.8 | 30.1 | 15.2 | 22.1 |
| 17001 | *Ltc4s* | 76 | 80.7 | 47.9 | 44.2 | 96.2 | 59.6 | 49.5 | 23.1 |
| 17436 | *Me1* | 116.7 | 118.8 | 76.2 | 62.6 | 122.1 | 110.9 | 56.6 | 61.7 |
| 68393 | *Mogat1* | 19 | 17.4 | 11.9 | 10.6 | 12.5 | 11.6 | 5.9 | 5.7 |
| 17850 | *Mut* | 40.2 | 34.7 | 24.2 | 19.8 | 34.9 | 30.6 | 15.8 | 19.4 |
| 107221 | *O3far1* | 22 | 18.7 | 8.3 | 9 | 12.8 | 12.7 | 6.6 | 4.1 |
| 13603 | *Opn3* | 19.4 | 17.4 | 7.8 | 9.8 | 11.8 | 12 | 3.4 | 7 |
| 74769 | *Pik3cb* | 13.7 | 12.5 | 7.4 | 7.4 | 13.2 | 10.8 | 6.5 | 5 |
| 116939 | *Pnpla3* | 45.5 | 67 | 25.3 | 16.5 | 39.7 | 62.4 | 24.5 | 15.4 |
| 99011 | *Pomt1* | 11 | 8.5 | 5.5 | 4.5 | 7.4 | 7.5 | 3.6 | 2.9 |
| 216974 | *Proca1* | 15.3 | 16.6 | 7.7 | 8.9 | 14.3 | 13.8 | 6.4 | 5.1 |
| 19139 | *Prps1* | 42.4 | 37.9 | 26.2 | 23.2 | 47.2 | 40 | 17.5 | 22.7 |
| 65116 | *Prrg2* | 10 | 10.9 | 6.5 | 4.8 | 10.5 | 9.4 | 5.7 | 4.4 |
| 19416 | *Rasd1* | 65.4 | 53.4 | 38 | 20 | 73.6 | 37.9 | 32.4 | 13.4 |
| 245945 | *Rbm47* | 10.2 | 10.9 | 8.4 | 5.8 | 6 | 7.5 | 2.9 | 3 |
| 71973 | *Rbpms2* | 16.8 | 18.2 | 10.1 | 9.6 | 14.2 | 12 | 5.9 | 5.2 |
| 57264 | *Retn* | 1387.5 | 1407.7 | 456.4 | 373.9 | 986.2 | 705.8 | 334.7 | 151.3 |
| 20503 | *Slc16a7* | 13.9 | 12.8 | 5.6 | 4.7 | 8.4 | 11.5 | 1.7 | 4.5 |
| 13358 | *Slc25a1* | 164.8 | 135.1 | 75.8 | 61.2 | 171.4 | 139.7 | 67.3 | 47.7 |
| 53896 | *Slc7a10* | 96.7 | 99.3 | 43 | 65.1 | 85.2 | 58.9 | 31.4 | 31.1 |
| 20661 | *Sort1* | 92.5 | 93 | 46.3 | 51.4 | 48.6 | 54.6 | 24.3 | 28.6 |
| 68792 | *Srpx2* | 9.7 | 13.2 | 4.8 | 4.8 | 6.9 | 11.1 | 3.5 | 3.7 |
| 20977 | *Syp* | 8.7 | 14.6 | 4.4 | 6 | 9.8 | 12.9 | 5.6 | 5.2 |
| 68385 | *Tlcd1* | 17.1 | 18.7 | 10.2 | 8.5 | 17.9 | 15.6 | 7.5 | 5.8 |
| 230157 | *Tmeff1* | 14.6 | 11.9 | 8.5 | 5.7 | 12.2 | 11 | 5.4 | 5 |
| 74122 | *Tmem43* | 134.7 | 126.3 | 64.1 | 85.1 | 80.4 | 77.4 | 39.7 | 40.1 |
| 22117 | *Tst* | 31 | 20.9 | 14.8 | 10.7 | 43 | 25.3 | 11.7 | 11.9 |
| 107448 | *Unc5a* | 6.1 | 6.2 | 3.8 | 4 | 5.1 | 4.8 | 3 | 2.5 |

**Supplemental Table 2: qRT-PCR primers**

| Gene | Forward primer 5'-3' | Reverse primer 5'-3' |
|---|---|---|
| *18s* | AGTCCCTGCCCTTTGTACACA | CGATCCGAGGGCCTCACTA |
| *Hira* | TTGACTTGGGATCCCGTTGG | AGTCTCTAGCTGCCAGTCCA |
| *Pparg* | TCGCTGATGCACTGCCTATG | GAGAGGTCCACAGAGCTGATT |
| *Cebpa* | CAAGAACAGCAACGAGTACCG | GTCACTGGTCAACTCCAGCAC |
| *Adipoq* | TGTTCCTCTTAATCCTGCCCA | CCAACCTGCACAAGTTCCCTT |
| *Glut4* | GTGACTGGAACACTGGTCCTA | CCAGCCACGTTGCATTGTAG |
| *Insig1* | TGTCGGTTTACTGTATCCCTGT | GTTGATGCCAACGAACACGG |
| *Retn* | AAGAACCTTTCATTTCCCCTCCT | GTCCAGCAATTTAAGCCAATGTT |
| *Slc25a1* | CAGAAGCAGTGGTAGTCGTG | TTGAGCCCTGCTTCAGTACA |
| *Glut9* | TTGCTTTAGCTTCCCTGATGTG | GAGAGGTTGTACCCGTAGAGG |
| *Glut2* | TCAGAAGACAAGATCACCGGA | GCTGGTGTGACTGTAAGTGGG |
| *Glut8* | AGCAGCGTCATGGAGATGC | ACAACGGTCAGTGTGAATAGGA |
| *Glut1* | CAGTTCGGCTATAACACTGGTG | GCCCCCGACAGAGAAGATG |
| *Fasn* | TTGACGGCTCACACACCTAC | TGTTCCACATCAAGAAACTGC |
| *Agpat2* | CAGCCAGGTTCTACGCCAAG | TGATGCTCATGTTATCCACGGT |
| *Bscl2* | AGACATGCAGAGACCAGATCAAA | GCCTCAGGCTTTGAGTTCCTG |
| *Cd36* | ATGGGCTGTGATCGGAACTG | TGTCTGTACACAGTGGTGCC |
| *Fatp1* | CGCCGATGTGCTCTATGACT | ACACAGTCATCCCAGAAGCG |
| *Chrebp* | AGATGGAGAACCGACGTATCA | ACTGAGCGTGCTGACAAGTC |

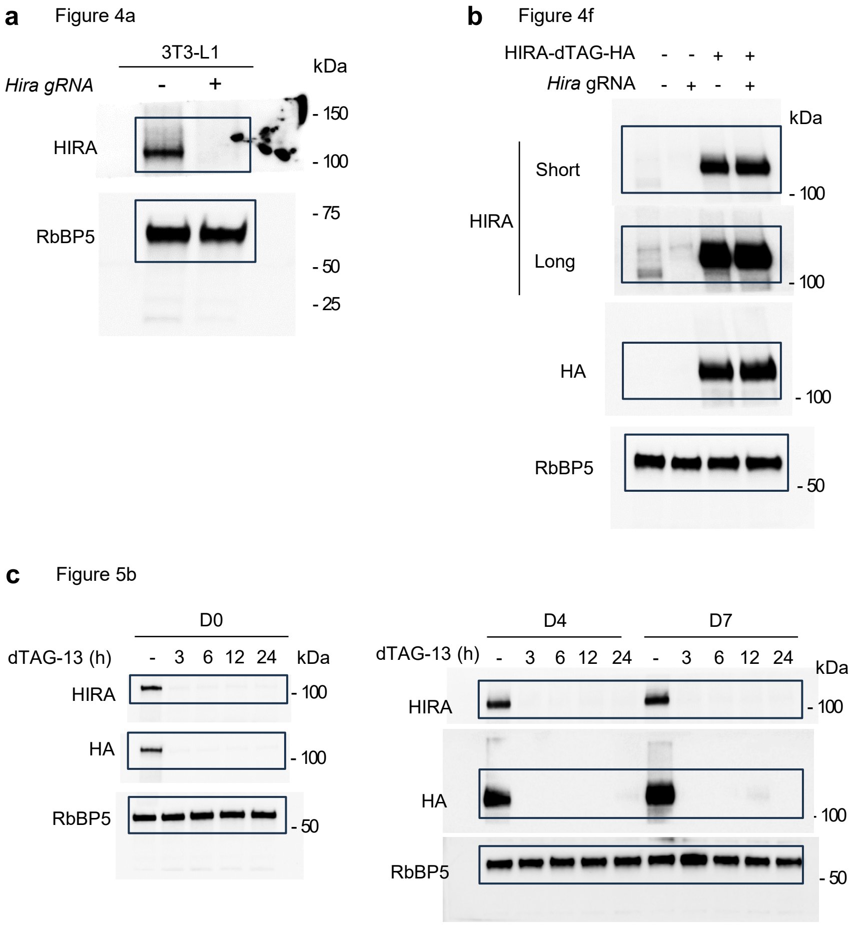


**Figure S11. Unprocessed WB images**
